# Positively twisted: The complex evolutionary history of Reverse Gyrase suggests a non-hyperthermophilic Last Universal Common Ancestor

**DOI:** 10.1101/524215

**Authors:** Ryan Catchpole, Patrick Forterre

**Affiliations:** Institut Pasteur, Unité de Biologie Moléculaire du Gène chez les Extrêmophiles (BMGE), Département de Microbiologie F-75015 Paris, France; Institute for Integrative Biology of the Cell (I2BC), CEA, CNRS, Univ. Paris-Sud, Univ. Paris-Saclay, 91198, Gif-sur-Yvette Cedex, France

## Abstract

Reverse gyrase (RG) is the only protein found ubiquitously in hyperthermophilic organisms, but absent from mesophiles. As such, its simple presence or absence allows us to deduce information about the optimal growth temperature of long-extinct organisms, even as far as the last universal common ancestor of extant life (LUCA). The growth environment and gene content of the LUCA has long been a source of debate in which RG often features. In an attempt to settle this debate, we carried out an exhaustive search for RG proteins, generating the largest RG dataset to date. Comprising 376 sequences, our dataset allows for phylogenetic reconstructions of RG with unprecedented size and detail. These RG phylogenies are strikingly different from those of known LUCA-encoded proteins, even when using the same set of species. Unlike LUCA-encoded proteins, RG does not form monophyletic archaeal and bacterial clades, suggesting RG emergence after the formation of these domains, and/or significant horizontal gene transfer. Even more strikingly, the branch lengths separating archaeal and bacterial groups are very short, inconsistent with the tempo of evolution from the time of the LUCA. Despite this, phylogenies limited to archaeal RG resolve most archaeal phyla, suggesting predominantly vertical evolution since the time of the last archaeal ancestor. In contrast, bacterial RG indicates emergence after the last bacterial ancestor followed by significant horizontal transfer. Taken together, these results suggest a non-hyperthermophilic LUCA and bacterial ancestor, with hyperthermophily emerging early in the evolution of the archaeal and bacterial domains.

## Introduction

Understanding the nature of the last universal common ancestor of extant life (LUCA) is one of the most difficult, yet important problems in evolutionary biology. If we were able to determine the genes encoded by the LUCA, we could make important conclusions regarding the evolutionary trajectories of all living organisms, as well as make predictions about the environment in which the LUCA lived. However, deciphering phylogenetic relationships dating back billions of years is a process fraught with difficulty. Not least because continual mutation over such time periods saturates sequences, erasing earlier phylogenetic signals that may exist, but also because mechanisms such as horizontal gene transfer act to blur phylogenies, further decreasing our ability to resolve such ancient relationships. Hence, the field of early evolutionary biology is one which is prone to disagreements, even when considering similar datasets. A poignant example of such disagreement comes from the phylogenies of Reverse Gyrase (RG), the only known hyperthermophile-specific protein, ubiquitously encoded by the genomes of hyperthermophilic organisms and absent from mesophiles (1–4). Understanding the evolutionary history of RG is important as the presence or absence of this gene in ancestral genomes (such as the LUCA) would allow us to infer a crude optimal growth temperature for these long-extinct species. The presence of reverse gyrase appears to be incompatible with mesophily, and, conversely, the absence of reverse gyrase appears incompatible with hyperthermophily. Thus, the presence of a gene encoding RG would infer a hyperthermophilic or thermophilic lifestyle excluding the option of a mesophilic lifestyle, and the absence, a mesophilic or moderately thermophilic growth condition, to the exclusion of hyperthermophily. This predictive ability is a powerful tool in evolutionary biology, where the optimal growth temperature of long-extinct organisms plays an important role in understanding genome evolution (e.g. “thermoreduction” (5)); protein and RNA evolution (6, 7) etc. In order to make inferences about the presence or absence of reverse gyrase in ancestral organisms, it is therefore vital to have a robust phylogeny for RG. However, the limited genetic data for hyperthermophilic organisms has restricted our ability to make such generalisations, and the small body of literature regarding RG evolution seems to flip-flop between the presence and absence of RG in LUCA.

Early analyses for the presence of RG were carried out experimentally by looking for positive supercoiling activity in cell lysates (8, 9). Though this allowed the identification of new RG-encoding species, these early experiments failed to detect RG in bacteria (and some archaeal species) due to the presence of the antagonistic DNA gyrase (10). Later, the discovery of RG activity in bacteria (namely in the *Thermotogales*) suggested that RG may be a more ancestral protein than previously thought, potentially evolving before the divergence of the bacterial and archaeal lineages (11). Even at this early stage in the absence of sequence data, the presence/absence of RG in the LUCA became an important factor in unravelling the nature of the ancestral life on earth (12). It wasn’t until sequence data for RG started to accumulate that phylogenetic analyses could be carried out. The first published phylogeny for RG included 13 sequences, and did not resolve the monophyly of the bacterial and archaeal domains (13). This result suggested that RG could not have evolved solely by vertical descent in the two domains, casting doubt over its presence in the LUCA. A later analysis *was* able to recover the bacterial and archaeal monophyly by using a dataset of 32 RG sequences; however, even then the domain separation was only weakly supported, and the bacterial tree did not reflect a canonical 16S phylogeny leading the authors to hypothesise an Archaea-to-Bacteria transfer for RG (4). Subsequently, another group was again able to recover the bacterial-archaeal domain separation in a phylogeny of only 15 sequences (2). Although these monophyly-recovering analyses suggest the direct descendance of RG from the LUCA, the datasets used were small and the intra-domain tree topologies were not as would be expected from an ancient protein evolving independently in the two domains. The most complete RG phylogeny to date formed part of a larger study searching for proteins present in the LUCA. Here, a dataset of 97 sequences identified RG as a candidate LUCA protein due to the monophyly of the bacterial and archaeal domains recovered in their analysis, prompting the authors to conclude that the LUCA was likely a hyperthermophile (14). Unfortunately, the topology of the recovered tree was not analysed in-depth and upon closer examination it is clear that this phylogeny suffers the same problems observed previously, that is, the branch between Archaea and Bacteria was rather short and the clades produced in the analysis are atypical and do not conform to the canonical 16S or universal protein phylogenies (tree reproduced in SI Fig. 1). Therefore, the conclusion that RG was encoded by the LUCA is supported weakly, at best.

With the quantity of genetic data increasing exponentially, and significant effort being made to sequence the genomes of archaeal species (many of which are hyperthermophiles), we thought it important to update the phylogeny of RG, and the evolutionary conclusions this can achieve. Using bioinformatics techniques, we reveal 376 RG sequences from 247 organisms across the bacterial and archaeal domains. Phylogenetic reconstruction of these sequences does not resolve the monophyly of the two domains, but rather reveals multiple potential horizontal transfer events. These results suggest RG was not present in the LUCA, but rather evolved after the divergence of the lineages leading to the LBCA and LACA. We therefore conclude that LUCA was a mesophile or moderate thermophile, with hyperthermophily evolving later, possibly before the emergence of the LACA.

## Materials and Methods

### Generation of Reverse Gyrase dataset

The 19 Reverse Gyrase sequences available in the Swiss-Prot database (15) were downloaded (July 2018) and aligned using MSAProbs v0.9.7 (16). The alignment was used to build a hidden Markov model (HMM) representative of confirmed RG proteins, using HMMER v3.1b2 (17) which was subsequently used as a query for a HMM search against the non-redundant protein database (downloaded 17 July 2018).

Hits were limited by a strict E-value cutoff of 10^−100^, and then aligned to identify hits which encode both a helicase and topoisomerase domain in a single amino-acid sequence (as per all RG sequences (18)). Alignments were viewed in Geneious 11.0.4 (https://www.geneious.com). A dataset of 371 putative RG sequences was recovered. A second search iteration (using all 371 sequences in generation of the query HMM) did not reveal any new RG sequences, and recovered the entire RG dataset.

### Split RG sequences

Known split RG sequences had to be added to the dataset manually as concatenations. To confirm the nature of split RG sequences, we used the entire output of the HMMer search to generate a simple phylogeny - alignment with ClustalW (19) and tree construction with Fasttree v2.1.9 (20), both performed on Galaxy@Pasteur (21) - to separate RG-encoding sequences from those of topoisomerase and helicase sequences. Sequences present in this RG clade, but excluded by our alignment-based hit-refining step were extracted from the tree, and themselves aligned with the 19 Swissprot RG sequences. These sequences indeed included the split RG sequences of the Nanoarchaeota and *Methanopyrus* species as well as truncated sequences (e.g. helicase-domain fragments of *Thermotoga maritima* RG used in structural analyses – 3OIY, 3P4Y, 3P4X), partial RG sequences recovered from metagenomic studies (e.g. KJR71718 from *Vulcanisaeta sp*. AZ3 and PSO07942 from Candidatus *Marsarchaeota* G2), and potential pseudogenisation and/or sequencing errors (e.g. WP_082398367 and WP_082398368 from *Aeropyrum camini* are encoded by two adjacent ORFs overlapping by 4bp which are out of frame by a single base). Potential new split RG sequences were also recovered in this analysis through visualisation of alignments in Geneious 11.0.4.

### RG sequence and species analyses

Sequence logos were generated using WebLogo 3.6.0 (22), and structural conservation mapping carried out with ConSurf 2016 (23).

Growth temperatures of RG-encoding organisms were obtained from BacDive (24) when possible, otherwise original research papers were sourced.

### Phylogenetic tree construction

Complete sequences corresponding to these Reverse Gyrase hits were downloaded, and again aligned with MSAProbs. Phylogenetically informative regions were selected using BMGE v1.12 (25), substitution model selected with ModelFinder (26) and phylogenetic trees were generated using IQ-TREE v1.6.6 (27). Bootstrap analysis was performed using ultrafast bootstrap approximation (1,000 replicates). Confirmation of trees was performed with RAxML v8.2.11 (28), and MrBayes v3.2.6 (29). Trees were visualised with iTol 4.2.3 (30).

Where phylogenies have concentrated on specific groups and/or particular groups of sequences have been removed from phylogenies, the reduced datasets were re-aligned, gaps removed, substitution model selected, and trees re-generated.

## Results

### Generation of Reverse Gyrase dataset

We screened the non-redundant protein database (downloaded 17 July 2018) for reverse gyrase sequences using an HMM built from the 19 Reverse Gyrase sequences available in the Swiss-Prot database.

The presence of helicase-like and topoisomerase-like sequences in our RG HMM (RG is a fusion between a SF2-like helicase domain and a Topoisomerase 1A domain (18)) resulted in the overwhelming presence of helicase- and topoisomerase-domain containing proteins in our search results, only a subset of which are RG sequences. Thus, hits were limited to those sequences which encode both a helicase and topoisomerase domain within a single amino acid sequence, representing 371 sequences in our dataset. Known RG sequences encoded by split genes e.g. those of *Methanopyrus kandleri* (31) and *Nanoarchaeum equitans* (32) were removed by this process and thus it was necessary to manually re-added these sequences to the dataset in a concatenated form taking the dataset to a total of 376 sequences. We were able to confirm the nature of these as split RG sequences (rather than distinct helicase and topoisomerase proteins) by their phylogenetic relatedness to known RG sequences (SI Fig. 2).

Interestingly, these analyses also revealed several potential split RG sequences previously unidentified e.g. protein pairs from *Aeropyrum camini*, Candidatus *Kryptonium thompsoni*, *Desulfurococcales* archaeon ex4484_217, and *Nitrososphaera* sp. (SI Fig. 3). However, of these sequences only the ORFs encoded by *Desulfurococcales* archaeon ex4484_217 are non-overlapping, thus, while recent gene splits cannot be ruled out, it is likely that the other pairs may be the result of sequencing errors.

Biochemical data will be necessary to elucidate whether potential split RG sequences code for functional enzymes. Due to the unknown nature of these sequences they were not added to our RG dataset, thus the final RG dataset contains 376 sequences.

### Analysis of RG sequences

The 376 putative RG sequences originate from 247 unique species (with intra-species variant sequences potentially arising from gene duplications and/or differences in start site annotation etc.). Alignment of the amino acid sequences reveals an average pairwise sequence identity of 34.2% across the entire dataset, with known helicase and topoisomerase motifs well conserved (SI Fig. 4). Mapping the degree of sequence conservation observed at each position in our multiple sequence alignment onto the structure of the RG protein from *Thermotoga maritima* pdb:4DDT (33) (identical results obtained with mapping to the RG structure from the archaeon *Archaeoglobus fulgidus* - pdb:1GKU (34)) reveals a high level of conservation within both the helicase and topoisomerase domains, with less conservation observed at exterior protein regions (Fig. 1a). Though this result is not hugely surprising considering the requirement of both domain activities for RG activity, it does suggest that our dataset is likely composed of true reverse gyrase proteins.

**Fig. 1.**
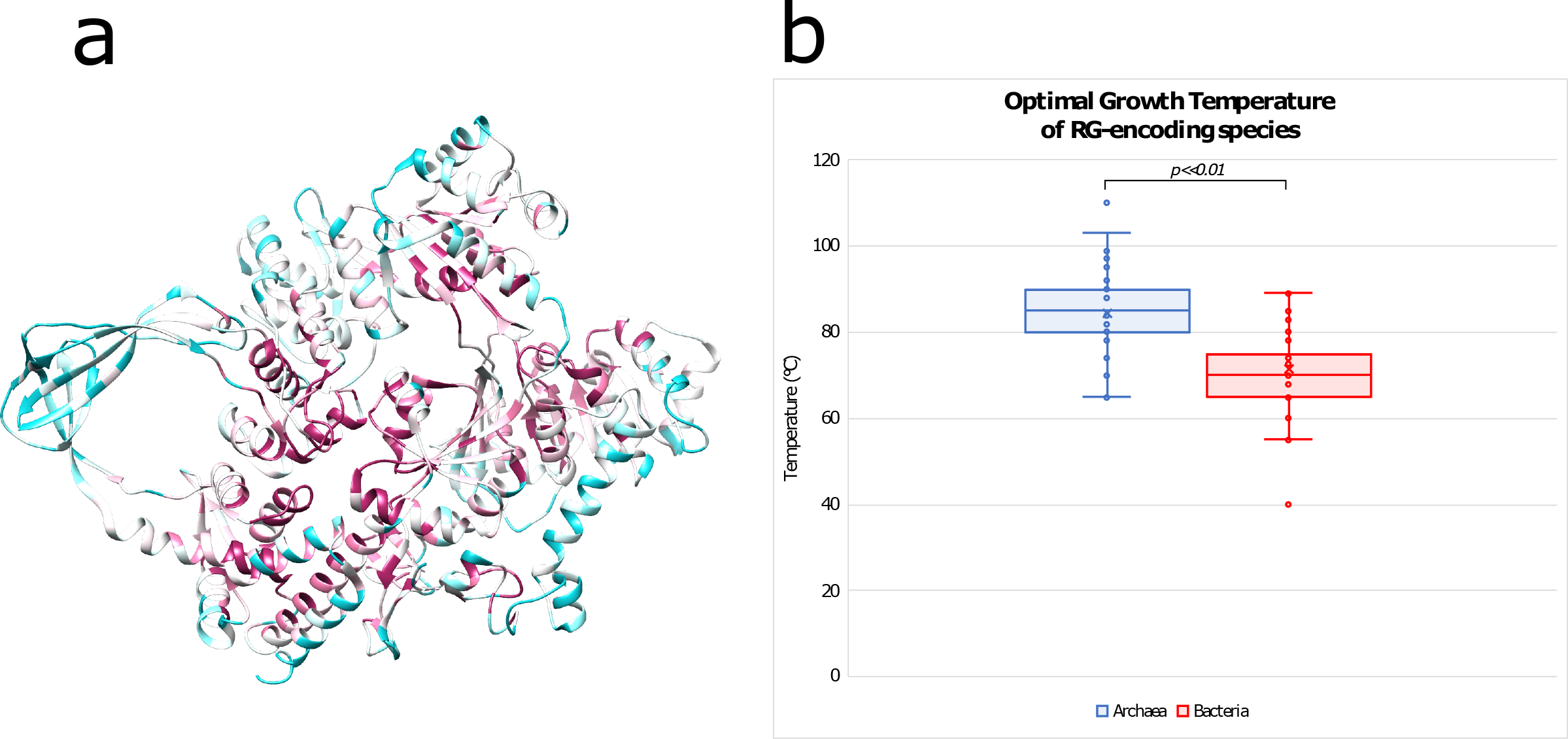
a) Sequence conservation recovered from alignment of RG dataset mapped onto RG structure from Thermotoga maritima (pdb:4DDT). Conserved residues indicated in red shades, less conserved residues indicated in blue shades. b) Optimal growth temperatures of RG-encoding species, separated by domain (Archaea in blue or Bacteria in red). RG-encoding Archaea have a significantly high mean growth temperature than RG-encoding bacteria.

### Species encoding RG

We were able to obtain information on the optimum growth temperatures of 174 of the 247 species encoding RG. As observed previously, almost all organisms encoding RG are hyperthermophiles or extreme-thermophiles, with 60% of the species in our dataset having an optimum growth temperature above 75°C and 89% having an optimum growth temperature above 65°C. Although difficult to confirm, we believe this dataset includes all hyperthermophilic organisms for which genome sequences are available. Thus, our data reaffirm the previous observation that RG is encoded by the genomes of all hyperthermophiles (1).

It is interesting to note that the archaeal species encoding RG tend to have significantly higher optimal growth temperatures than the bacterial species (Fig. 1b). The average optimal growth temperature for RG-encoding archaea is 84.3°C, whereas it is only 71.4°C for bacteria. This is further reinforced by the fact that 83% of the archaeal species with RG have optimum growth temperatures above 75°C, in contrast with only 21% of bacterial species.

In addition to extreme thermophiles, our search also gave hits to RG sequences in 5 moderate thermophiles with optimum growth temperatures below 65°C: *Thermodesulfovibrio aggregans* (60°C (35)); *Nitratiruptor tergarcus* (55°C (36)); *Lebetimonas natsushimae* (55°C (37)); *Caminibacter mediatlanticus* (55°C (38)); *Nautilia profundicola* (40°C (39)). The presence of RG in *N. profundicola* has been described previously and is likely an adaptation to short-term exposure to elevated temperatures in hydrothermal vent environments, with RG expression increasing 100-fold during temperature stress at 65°C (40). As *Nitratiruptor tergarcus, Lebetimonas natsushimae* and *Caminibacter mediatlanticus* were also isolated from the walls of active hydrothermal vents, similar adaptive mechanisms may explain the presence of RG in these species.

### Phylogenetic analysis of RG dataset

In order to investigate the evolutionary history of the RG protein as a whole, we used our entire RG dataset to generate a single phylogenetic tree. Due to the presence of non-informative residues in our alignment (e.g. an intein, difference in *in silico* start site prediction, poorly conserved residues etc), it was important to process the dataset to remove such positions. Trimming the dataset with BMGE resulted in an alignment of 571 positions. We were initially concerned that such severe trimming could remove important data from the alignment and decrease branch lengths in resulting phylogenies (41), thus the analyses were repeated with only light trimming (removing only 17% of positions) using Noisy (42). The resulting tree resolved similar clades with similar relative branch lengths to that generated with BMGE (SI Fig. 5 vs SI Fig. 6).

The first feature that is clear to note in our complete RG phylogeny is that the bacteria and archaeal sequences are not monophyletic (Fig. 2). The bacterial sequences split into 3 different clades, as do the archaeal sequences.

**Fig. 2.**
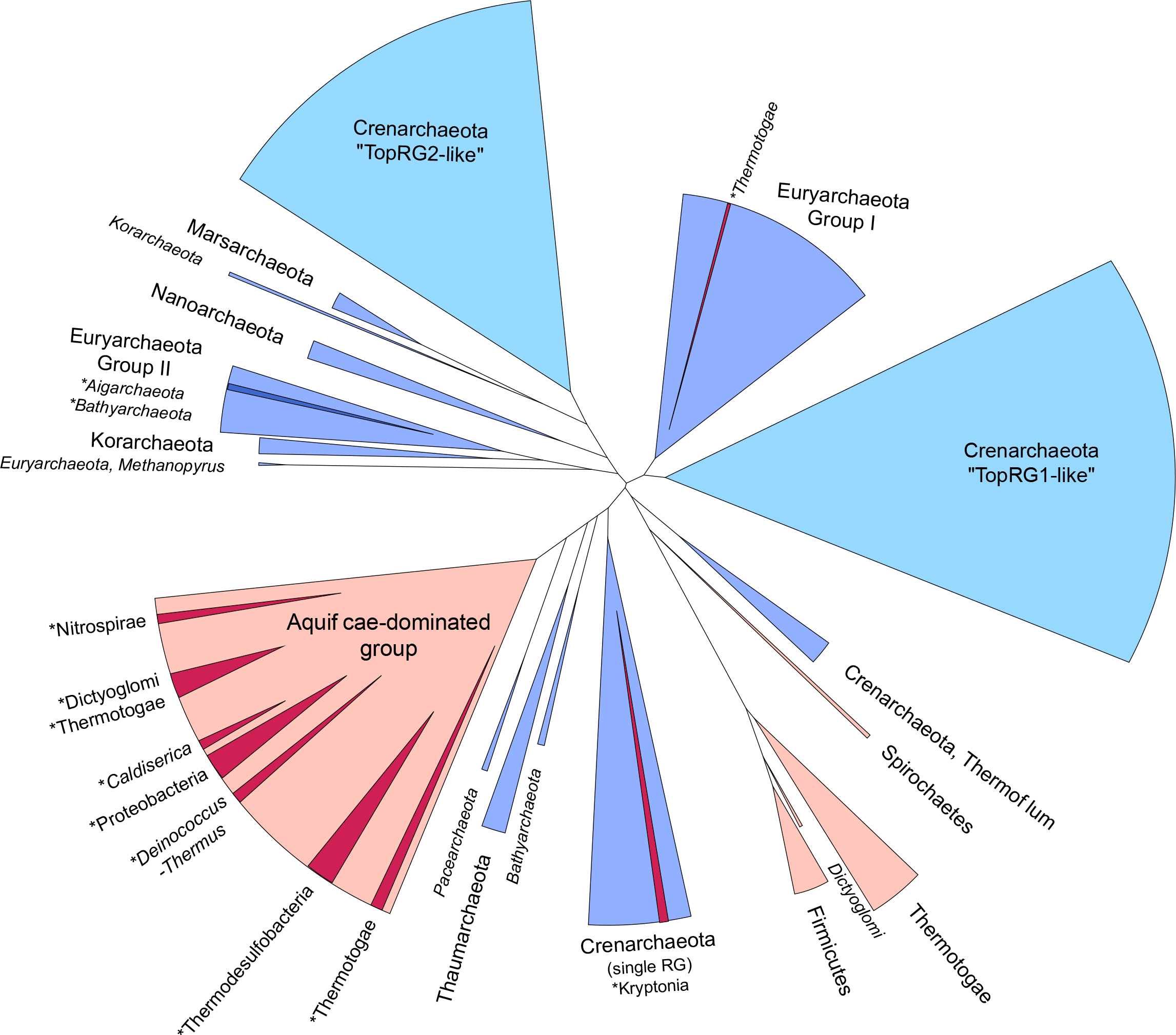
Schematic representation of phylogenetic tree generated using entire RG dataset. Archaeal clades coloured in blue, Bacterial clades in red, with phyla indicated. Clades formed inside canonical phyla are indicated in darker shades, and labelled with an asterisk. Clades labelled with italicised text indicate ≤2 sequences present. Crenarchaeal TopRG1-like and TopRG2-like paralogues indicated in pale blue. Detailed tree available in SI Fig. 6.

This is in direct contrast to that which would be expected if RG had emerged in a common ancestor of bacteria and archaea (i.e. the LUCA), and evolved independently since the divergence of the bacterial and archaeal lineages. In order to illustrate this contrast, we collected the sequences of two universal marker proteins (rpoB encoding the RNA polymerase β-subunit, and EF-G/aEF-2 encoding translation elongation factor G), as well as the 16S rDNA from the RG-encoding species recovered in our original RG search. These datasets were used to generate phylogenetic trees in a manner identical to that used for the RG dataset. In contrast to the RG phylogeny, these datasets show a clear monophyletic separation of the bacterial and archaeal species (Fig. 3, 16S in SI Fig. 7). This is exactly as would be expected for sequences which diverged before the appearance of the LACA and the LBCA (i.e. sequences present in the LUCA).

**Fig. 3.**
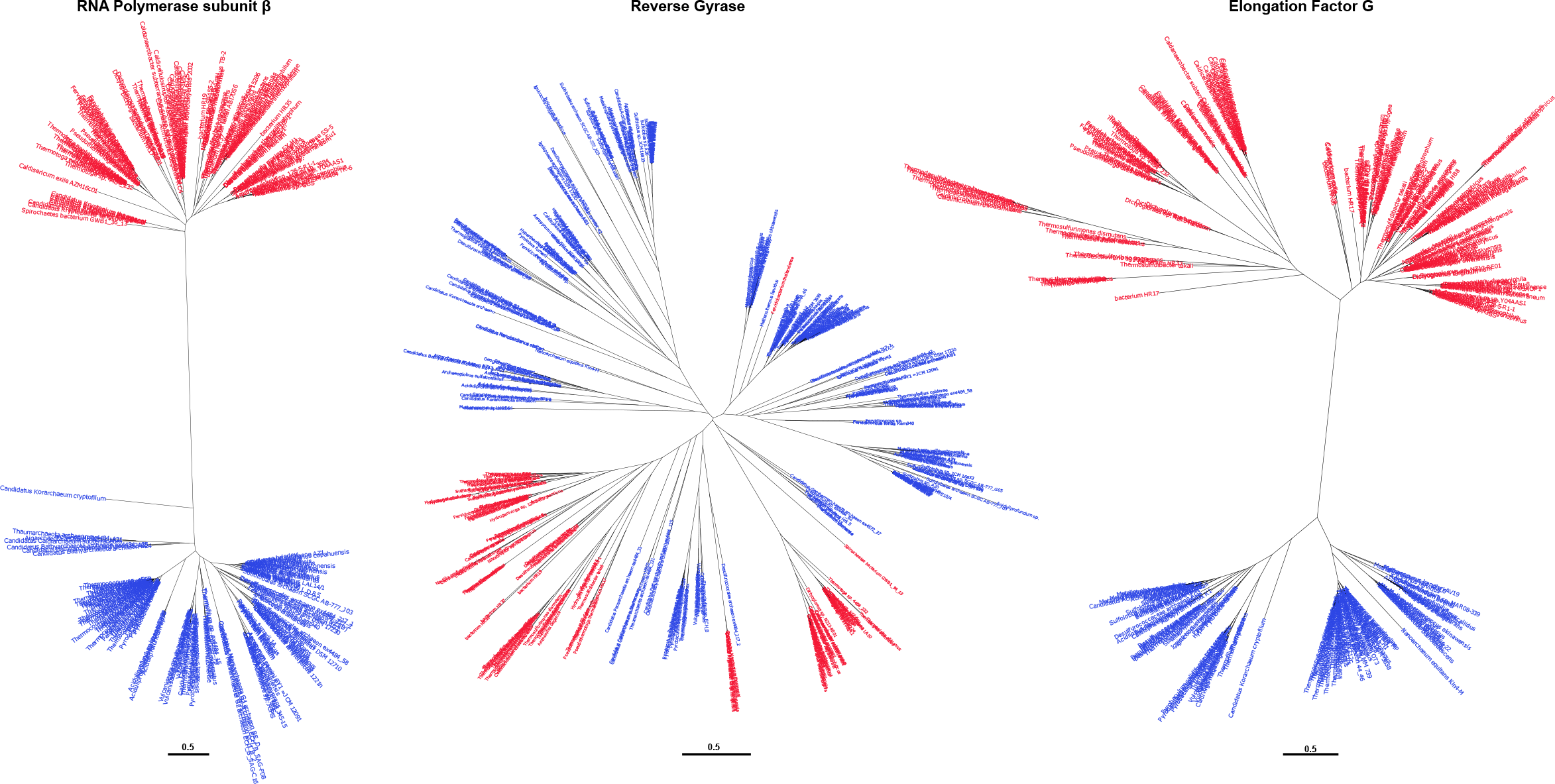
Phylogenetic trees generated using universal proteins from RG-encoding species, compared with RG itself. First panel, RNA Polymerase subunit β; second panel, reverse gyrase; third panel, Elongation Factor G. In all trees, sequences encoded by Archaeal species are indicated in blue, Bacterial species in red.

Perhaps more strikingly, the lengths of branches separating the bacterial and archaeal species are clearly different between our RG phylogeny and those of our universal marker protein phylogenies (Fig. 3, SI Fig. 7). The very long branch lengths displayed by our universal marker proteins are in agreement with the idea that the tempo of evolution was much higher during the period between LUCA and the specific ancestors of Archaea and Bacteria, decreasing later on during the diversification of these two domains (43, 44). This results in the formation of two very divergent versions of universal proteins in Bacteria and Archaea, separated in phylogenetic trees by a very long branch. In contrast, the RG phylogenetic tree has much shorter inter-node branch lengths, inconsistent with such an evolutionary scenario. This result indicates that archaeal and bacterial RG do not form two distinct versions, and thus likely diverged from each other when the tempo of evolution had already slowed down i.e. shortly before or after the formation of the two prokaryotic domains. When combined with the polyphyletic nature of the bacterial and archaeal RG sequences, it becomes clear that RG must have evolved after the time of the LUCA. This result is in agreement with that observed in earlier RG phylogenies (4, 13), but contrasts with more recent reconstructions (2, 14).

### Crenarchaeal RG duplication

An oddity in RG sequences is observed in a large group of Crenarchaeota, including the well-studied Sulfolobales. Here, RG has undergone a gene duplication, and this duplication is clearly represented in our RG phylogenetic tree where the two paralogs form two distinct clades (Fig. 2). Additionally, it is apparent from *in vitro* and *in vivo* experiments that the two RG paralogs have diverged in function (45, 46), with non-complementary activities (and neither being essential in *Sulfolobus islandicus* (47). For simplicity, we refer to these RG paralogs by their nomenclature in *Sulfolobus* species i.e. TopR1-like and TopR2-like. TopR1 has been reported to function as a classical RG, exhibiting ATP-independent topoisomerase activity and DNA renaturation at high temperature; whereas the function of TopR2 is less clear, seemingly exhibiting high levels of ATP-dependent supercoiling at temperatures below those usually required for cell division (45). In order to analyse whether this divergence in activity was mirrored by changes in the RG amino-acid sequence, we used datasets limited to each paralog to generate alignments of TopR1-like and TopR2-like proteins. Comparison of these alignments with each other, and with that generated using the complete RG dataset reveals that all of the conserved motifs (both helicase and topoisomerase domain motifs) are present in both TopR1-like and TopR2-like sequences, as well as the active site tyrosine. Despite the similarities, we observed a notable difference in the second putative zinc finger motif of RG (Zn2) of the Crenarchaeal paralogs. The Zn2 motif is conserved in only 62% of our RG sequences, however, it is strictly conserved among all TopR2-like sequences. In contrast, only 35% (23/65) of TopR1-like sequences encode the second cysteine in this CxxCx_9-11_CxxC motif, with around half of those (11/23) containing additional inserts within Zn2. This motif has been shown to be important for DNA binding and positive supercoiling, but not for relaxation of negative supercoils or for ATPase activity (33, 48) and thus may explain the differences in processivity and function of these two enzymes.

Due to the apparent functional divergence of TopR1-like and TopR2-like proteins, as well as divergence of TopR2-like proteins from the canonical RG functionality, we chose to remove these sequences from our RG phylogeny. Not only could the different evolutionary trajectory of TopR1-like and TopR2-like proteins alter the tree structure (e.g. due to long-branch attraction artefacts), TopR2-like proteins do not seem necessary for growth at high temperature, and thus are not informative as to the hyperthermophily of ancestral species. Removal of these sequences did not resolve the monophyly of the bacterial and archaeal domains, nor did it increase the inter-domain branch lengths (Fig. 4, SI Fig. 8). Despite having minimal impact on tree topology and thus on our conclusions regarding RG presence in LUCA, we chose to carry out subsequent analyses without TopR1-like or TopR2-like proteins.

**Fig. 4.**
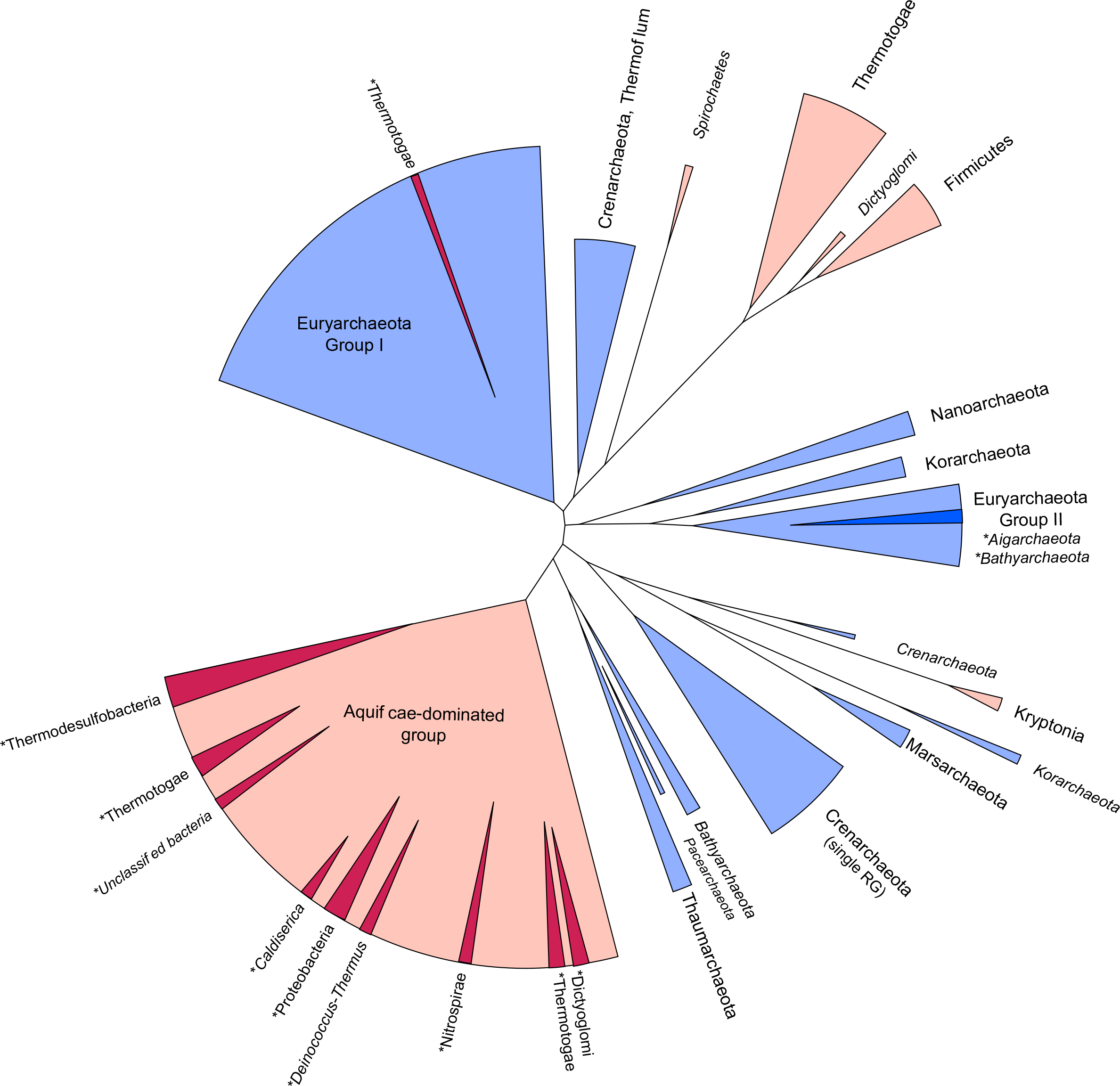
Schematic representation of phylogenetic tree generated using RG dataset without TopR1-like and TopR2-like sequences. Archaeal clades coloured in blue, Bacterial clades in red, with phyla indicated. Clades formed inside canonical phyla are indicated in darker shades, and labelled with an asterisk. Clades labelled with italicised text indicate ≤2 sequences present. Detailed tree available in SI Fig. 8.

### Domain-specific RG phylogenies

The absence of RG in LUCA suggests that the protein had to have emerged in either the lineage leading to the LACA (or in a more recent archaeal group), or in the lineage leading to the LBCA (or in a more recent bacterial group), with horizontal gene transfer spreading RG to the second domain. These two scenarios lead, *a priori*, to hypotheses testable by further phylogenetic analyses: if RG evolved in the lineage leading to the LACA, an RG phylogeny produced using only archaeal sequences should preserve the canonical archaeal taxonomic groups, whereas the inter-domain HGT required to introduce RG into the bacterial domain would likely not produce a typical bacterial taxonomy (and *vice versa*).

In order to test this hypothesis, we generated RG datasets containing only the 125 archaeal RG sequences (excluding TopR1-like and TopR2-like sequences), or containing only the 118 bacterial sequences (Fig. 5, SI Fig. 9, 10). These new datasets were then reanalysed in the same manner as the entire RG dataset.

**Fig. 5.**
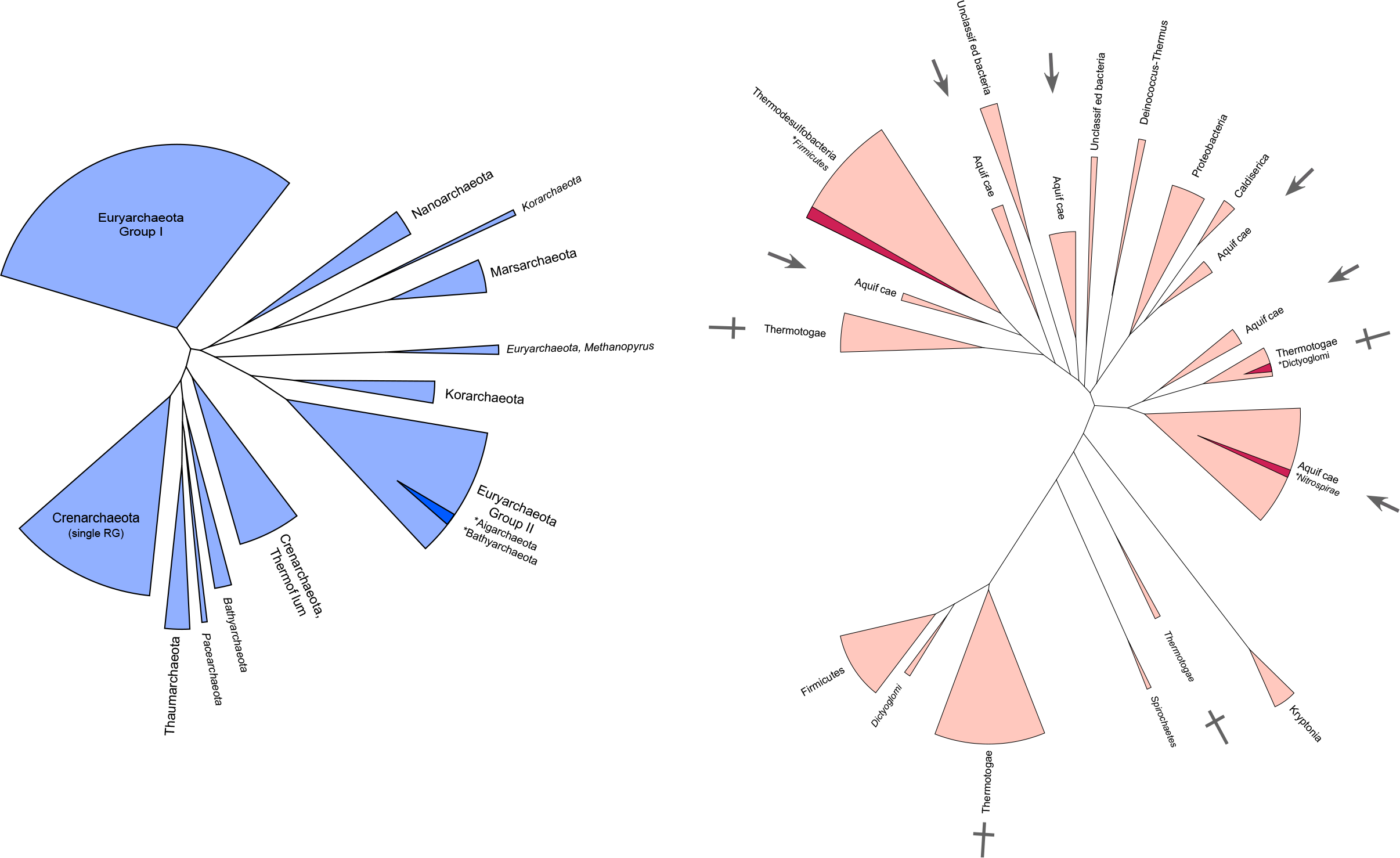
Schematic representation of phylogenetic trees generated using only Archaeal RG sequences (blue), or Bacterial RG sequences (red). Clades formed inside canonical phyla are indicated in darker shades, and labelled with an asterisk. Clades labelled with italicised text indicate ≤2 sequences present. Aquificae and Thermotoga clades are labelled with arrow and cross, respectively, to highlight their paraphyletic nature.

The archaeal RG phylogeny resulting from this analysis resolves into separate clades rather congruent with the consensus archaeal phylogeny but with several unexpected positions (Fig. 5). All group I Euryarchaeota (*Thermococcus*, *Methanococcus, Methanobacteriales*) group together, with the exception of *Methanopyrus kandleri* which is known to be fast evolving and difficult to position in the archaeal tree (49). *Methanopyrus kandleri* and three RG from Korarchaea (other fast evolving archaeal species) branch at the base of a clade grouping Euryarchaea group II (*Aciduliprofundum* and *Archaeoglobales*). A RG from a candidatus Bathyarchaeon branches within group II archaea, suggesting either contamination of this metagenome or lateral gene transfer from a group II euryarchaeon to a Bathyarchaeon. Two other bathyarchaeal RG branch with a Thaumarchaeon and two candidatus Caldiarchaeum subterraneum. This grouping is consistent with the monophyly of the BAT (Bathyarchaea, Aigarchaea, Thaumarchaea) clade observed in most phylogenetic analyses (50, 51). However, in the RG phylogeny, this BAT clade branches within Crenarchaeota, reminiscent of the TACK grouping recovered in previous phylogenies (52). Another unexpected position is that of the Marsarchaeota that usually branch as sister group of Crenarchaeota in phylogenetic analyses (53), and that branch within Euryarchaea in the RG tree, together with a single Korarchaeota and Nanoarchaea (also fast evolving species and thus difficult to place phylogenetically). It is unclear if these anomalies are due to some gene transfer events between Archaea (from Crenarchaea to archaea of the BAT group and from Euryarchaea to Marsarchaeota) or from the low resolution of the RG tree at the interphylum level (branch support for these spurious clades is weak). Intriguingly, the archaeal phylogeny is perhaps better resolved in RG trees where bacterial sequences are present – Fig. 4, a full RG tree (without TopRG1 or TopRG2-like sequences) reveals Marsarchaeota located in their canonical position close to the Crenarchaeota and the BAT group distinct from Crenarchaeota. These results are further difficult to interpret since there is no consensus to the position of the root of the archaeal tree (54, 55), and thus no ‘true’ phylogeny to which we can compare RG evolution. Regardless, the clear-cut separation between Crenarchaeota and Euryarchaeota (Fig. 2, 4; SI Fig. 1) suggest that RG was introduced into the archaeal domain before their divergence. It is unclear if RG was already present in the LACA, though the recovered divergence of the Thaumarchaeota and Bathyarchoaeta from the Crenarchaeota and Euryarchaeota might suggest this to be the case. More sequences from basal archaeal groups (e.g. Thaumarchaeota and Bathyarchaeota), as well as under-represented groups (e.g. Nanoarchaeota and Korarchaeota) will help to strengthen the archaeal RG tree, and allow more unambiguous extrapolation to the LACA.

Notably, the bacterial RG phylogeny follows a much more random pattern of clade separation than the archaeal one. This is exemplified by the Thermotogales and Aquificales, where members of these orders can be seen in 4 and 6 separate clades, respectively (Fig. 5). Moreover, the major clade of Aquificae and that of the Thermodesulfobacteria are separated by quite some distance, with many other clades branching between them. This result is very different from that observed previously with 16S phylogenies (SI Fig. 7), and also with our phylogenies based on universal proteins, where the Aquificae and Thermodesulfobacteria are closely related groups. Furthermore, the Thermotoga form a clade together with the Firmicutes – a branching which is not unreasonable; yet the very closely related Pseudothermotoga form a clade within an Aquificae sub-group. Similarly to the Archaea, no universally accepted Bacterial phylogeny is available, and thus comparing the order of bacterial clades resolved with RG is difficult. However, it is clear that the bacterial RG phylogeny does not even conform to canonical taxonomy, and thus is likely highly influenced by gene transfer events.

Taken together, these results suggest that RG has evolved mostly vertically in the Archaea before the divergence of Euryarchaea and Crenarchaea, partly preserving the evolutionary history of the Archaea within its sequence. The evolution of RG in the bacteria has not followed such a pattern of vertical inheritance, rather several horizontal transfer events have resulted in the movement of RG within the bacterial domain.

### Evidence for horizontal transfer of RG

The dataset of bacterial RG proteins shows further evidence of HGT at the level of individual species or genera. For example, some members of the Aquificae (including *Aquifex aeolicus* and *Hydrogenivirga sp. 128-5-R1-1*) encode two copies of RG proteins. In these cases, one of the RG copies is most closely related to other RG proteins encoded by Aquificales; however, the second RG appears most closely related to that of Thermodesulfobacteria (SI Fig. 8). If these two RG copies had arisen by duplication within an ancestor of *Aquifex* and *Hydrogenivirga*, we might expect the two proteins to be most closely related to each other rather than to RG of other species. Instead, it appears that the second RG copy has arisen by HGT from an ancestor of Thermodesulfobacteria. Alternatively, the two RG sequences could have arisen by duplication within an ancestor of Aquificales, and then each transferred independently to other organisms, thus disrupting the expected tree topology. Either way, it is clear that HGT has played a significant role in the evolution of RG in these groups.

Our RG phylogeny also confirms previously observed evidence for HGT of RG. For example, the inter-domain transfer from a Crenarchaeon to an ancestral Kryptonia bacterium (56), and the non-canonical position of some *Dictyoglomus* species suggesting transfer from a *Fervidobacterium* (57) (SI Fig. 8).

### Rooting of the RG phylogenetic tree

The rooting of the reverse gyrase tree could potentially bring new arguments in favour of specific scenarios for the origin of RG. If RG initially originated in Archaea and was present in this domain before the diversification of Euryarchaea and other Archaea, the tree should be *a priori* rooted in the archaeal domain. RG is thought to have arisen from a gene fusion event between a helicase and a topoisomerase 1A domain, thus we expect that both helicase- and topoisomerase 1A-containing proteins could act as an appropriate outgroup to root the archaeal tree. We selected the topoisomerase domain since its larger size made it a more appropriate candidate for rooting the archaeal tree. We used topoisomerase sequences with a known phylogenetic relationship taken from Forterre et al. 2007 (58). Including these sequences with our dataset of RG sequences (excluding the TopR1 and TopR2-like sequences) produces a tree with three well separated clades corresponding to RG, Bacterial Type IA DNA topoisomerases (the orthologues of the *Escherichia* omega-protein), and archaeal type IA DNA topoisomerases (also often annotated as DNA topoisomerase III). In this tree, the RG turned out to be rooted in one of the bacterial RG clades (SI Fig. 11). The same rooting was obtained with bacterial or archaeal Topo IA alone (SI Fig. 12), suggesting that RG may have emerged within an ancestral bacterial group. This bacterial group consists of Thermodesulfobacteria, Aquificae, and Thermotoga species; often represented as deeply branching bacteria (Colman et al. 2016). Interestingly, despite the rooting of RG within a bacterial clade, the archaeal phylogeny is still congruent with canonical archaeal trees. This result suggests that RG was indeed present very early in the history of archaea, possibly in LACA, but surprisingly also suggests that RG might have been also present very early in some Bacteria, and transferred to an ancestral archaeal lineage. It is worth noting that this rooting in a bacterial RG clade was lost when the Crenarchaeal TopR1-like proteins were included in the analyses (SI Fig. 13), with the RG root falling between a bacterial and archaeal clade. If either of these rootings were confirmed in future analyses, one could imagine evolutionary scenarios to explain them. For instance, RG could have originated in a subgroup of Bacteria (e.g. an ancestor of Thermodesulfobacteria) and later transferred to the archaeal lineage before the emergence of LACA (at the very least, before the divergence of the Crenarchaeota and Euryarchaeota, see above), giving a bacterial rooting. Alternatively if RG originated in the archaeal stem lineage (between the LUCA and the LACA), it could have been transferred from a member of this lineage to a subgroup of Bacteria before the LACA, giving a root between bacterial and archaeal clades. Notably, both of these scenarios would imply that the LBCA emerged before the LACA, a possibility which, to our knowledge, has not been considered up to now. However, considering the large distance between RG and the outgroup sequences, and the variability of rooting obtained depending upon the RG dataset selected, the above scenarios should be interpreted with much caution.

## Discussion

The work presented here strongly suggests the absence of RG in the LUCA since the archaeal and bacterial RG do not form two monophyletic clades in our phylogenetic analyses. Furthermore, the short branch length between any inter-domain clades of our phylogenies (Fig. 4) indicate a period of divergence inconsistent with the tempo of evolution between LUCA and the common ancestors of Archaea and Bacteria; the branches between the different archaeal and bacterial clades are all very short, suggesting the existence of a single version of RG. In contrast, using the same set of species, we have shown here that not only do Archaea and Bacteria form two monophyletic clades in phylogenies of markers known to be present in LUCA such as EF-G, RNA polymerase and 16S rRNA, but also that the branch between these two clades is very long (Fig. 3). Using a different set of species, we also systematically observed long branches separating Archaea and Bacteria in the phylogenies of 36 universal proteins (most likely present in the LUCA) except for a handful of very small ribosomal proteins (51). Our RG results are in contradiction with those of Weiss and colleagues who recently concluded that RG was present in LUCA in their tentative reconstruction of the LUCA proteome (14) (SI Fig. 1 vs SI Fig. 6). This could be explained by the difference in the number of sequences used in the two analyses (347 and 97, respectively) and the fact that Weiss and colleagues do not include the branch length between the Archaea and Bacteria as a criterium to conclude that a protein was present or not in the LUCA. This branch was also very short in the RG tree of Weiss and colleagues (SI Fig. 1) and we could not ourselves recover the monophyly of Archaea and Bacteria using their dataset and tree construction method, suggesting that the monophyly versus paraphyly of Archaea and Bacteria is sensitive to some parameters of tree reconstruction, further suggesting a non-distinct separation of these usually highly divergent domains.

The absence of RG in LUCA, combined with the apparent requirement of RG for growth at high temperature, suggests the existence of a non-hyperthermophilic LUCA. This is consistent with the work of Manolo Gouy and colleagues on ancestral protein and rRNA reconstructions (6, 7, 59). These authors suggested that the LUCA was either a mesophile or a moderate thermophile based on the rRNA GC percent and on the amino-acid composition of its proteome. A non-hyperthermophilic LUCA is also in agreement with the idea that LUCA was an organism simpler than modern ones, with smaller ribosomes (60) and possibly an RNA genome (61). The origin of most DNA replication proteins cannot be traced back to LUCA (62), and it seems that RG is not an exception. The few universal proteins that manipulate DNA in modern organisms could have been working on RNA in the LUCA or transferred independently from DNA viruses to the archaeal and bacterial lineages (63). RNA being extremely fragile at high temperature (64) a mesophilic LUCA fits also well with the hypothesis of LUCA having an RNA genome. The transition from a LUCA with an RNA genome to LACA and LBCA with DNA genomes could also explain why the tempo of evolution drastically slowed down between LUCA and the two prokaryotic ancestors, considering that DNA can be replicated and repaired much more faithfully than RNA (Forterre, 2006). With respect to our RG phylogenies, and RG evolution in general, the short branch lengths between bacterial and archaeal clades would place the emergence of RG in the age of DNA cells i.e. more recently than the time of a rapidly evolving RNA-based LUCA (and post-LUCA lineage). This, perhaps, would seem logical considering the strict DNA substrate-dependence of RG, and RG conferring adaptation to hyperthermophilic growth temperatures – a state likely incompatible with RNA genomes. Finally, our work suggest that the criteria of branch length should be considered, in addition to the monophyly of Archaea and Bacteria, in establishing which proteins were probably present or not in LUCA.

