## Supplementary material for "Positively twisted: The complex evolutionary history of Reverse Gyrase suggests a non-hyperthermophilic Last Universal Common Ancestor": SI Fig. 1

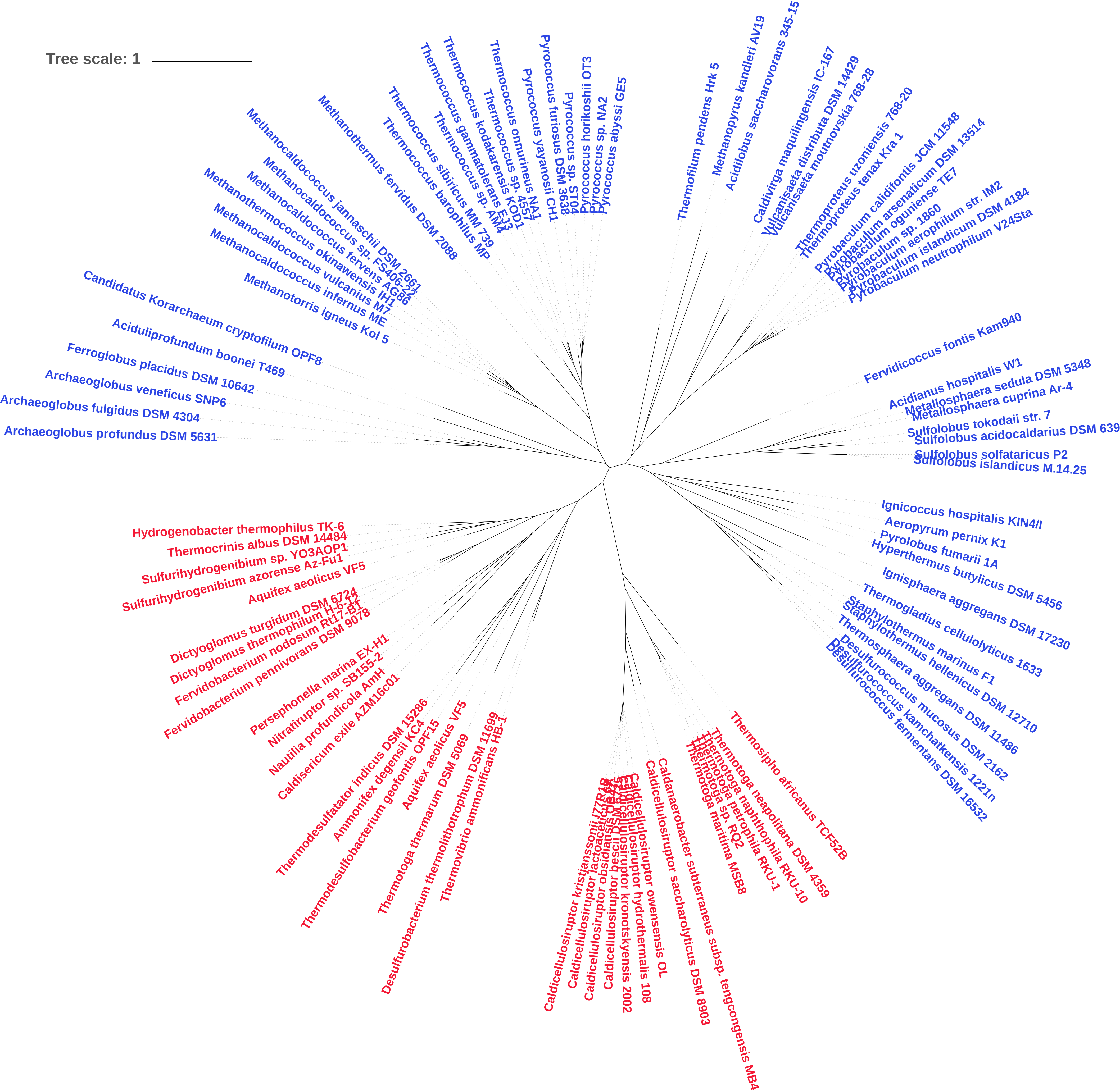
Tree scale: 1
Thermofilum pendens Hrk 5
Methanopyrus kandleri AV19
Acidilobus saccharovorans 345-15
Caldivirga maquilingensis IC-167
Vulcanisaeta distributa DSM 14429
Vulcanisaeta moutnovskia 768-28
Thermoproteus uzoniensis 768-20
Thermoproteus tenax Kra 1
Pyrobaculum calidifontis JCM 11548
Pyrobaculum arsenaticum DSM 13514
Pyrobaculum oguniense TE7
Pyrobaculum sp. 1860
Pyrobaculum aerophilum str. IM2
Pyrobaculum islandicum DSM 4184
Pyrobaculum neutrophilum V24Sta
Fervidicoccus fontis Kam940
Acidianus hospitalis W1
Metallosphaera sedula DSM 5348
Metallosphaera cuprina Ar-4
Sulfolobus tokodaii str. 7
Sulfolobus acidocaldarius DSM 639
Sulfolobus solfataricus P2
Sulfolobus islandicus M.14.25
Ignicoccus hospitalis KIN4/I
Aeropyrum pernix K1
Pyrolobus fumarii 1A
Hyperthermus butylicus DSM 5456
Ignisphaera aggregans DSM 17230
Thermogladius cellulolyticus 1633
Staphylothermus marinus F1
Staphylothermus hellenicus DSM 12710
Thermosphaera aggregans DSM 11486
Desulfurococcus mucosus DSM 2162
Desulfurococcus kamchatkensis 1221n
Desulfurococcus fermentans DSM 16532
Thermosipho africanus TCF52B
Thermotoga neapolitana DSM 4359
Thermotoga naphthophila RKU-10
Thermotoga petrophila RKU-1
Thermotoga sp. RQ2
Thermotoga maritima MSB8
Caldanaerobacter subterraneus subsp. tengcongensis MB4
Caldicellulosiruptor saccharolyticus DSM 8903
Caldicellulosiruptor owensensis OL
Caldicellulosiruptor hydrothermalis 108
Caldicellulosiruptor kronotskyensis 2002
Caldicellulosiruptor kristjanssonii I77R1B
Thermovibrio ammonificans HB-1
Desulfurobacterium thermolithotrophum DSM 11699
Thermotoga thermarum DSM 5069
Aquifex aeolicus VF5
Thermodesulfobacterium geofontis OPF15
Ammonifex degensii KC4
Thermodesulfatator indicus DSM 15286
Caldisericum exile AZM16c01
Nautilia profundicola AmH
Nitratiruptor sp. SB155-2
Persephonella marina EX-H1
Fervidobacterium pennivorans DSM 9078
Fervidobacterium nodosum Rt17-B1
Dictyoglomus thermophilum H-6-12
Dictyoglomus turgidum DSM 6724
Aquifex aeolicus VF5
Sulfurihydrogenibium azorense Az-Fu1
Sulfurihydrogenibium sp. YO3AOP1
Thermocrinis albus DSM 14484
Hydrogenobacter thermophilus TK-6
Archaeoglobus profundus DSM 5631
Archaeoglobus fulgidus DSM 4304
Archaeoglobus veneficus SNP6
Ferroglobus placidus DSM 10642
Aciduliprofundum boonei T469
Candidatus Korarchaeum cryptofilum OPF8
Methanotorris igneus Kol 5
Methanocaldococcus infernus ME
Methanocaldococcus vulcanius M7
Methanothermococcus okinawensis IH1
Methanocaldococcus fervens AG86
Methanocaldococcus sp. FS406-22
Methanocaldococcus jannaschii DSM 2661
Methanothermus fervidus DSM 2088
Thermococcus barophilus MP
Thermococcus sibiricus MM 739
Thermococcus sp. AM4
Thermococcus gammatolerans EJ3
Thermococcus kodakarensis KOD1
Thermococcus sp. 4557
Thermococcus onnurineus NA1
Pyrococcus yayanosii CH1
Pyrococcus furiosus DSM 3638
Pyrococcus sp. ST04
Pyrococcus horikoshii OT3
Pyrococcus sp. NA2
Pyrococcus abyssi GE5

**SI Fig. 1.** Reproduction of RG phylogeny constructed by Weiss et al. 2016 (1)
