## Supplementary material for "Positively twisted: The complex evolutionary history of Reverse Gyrase suggests a non-hyperthermophilic Last Universal Common Ancestor": SI Fig. 2

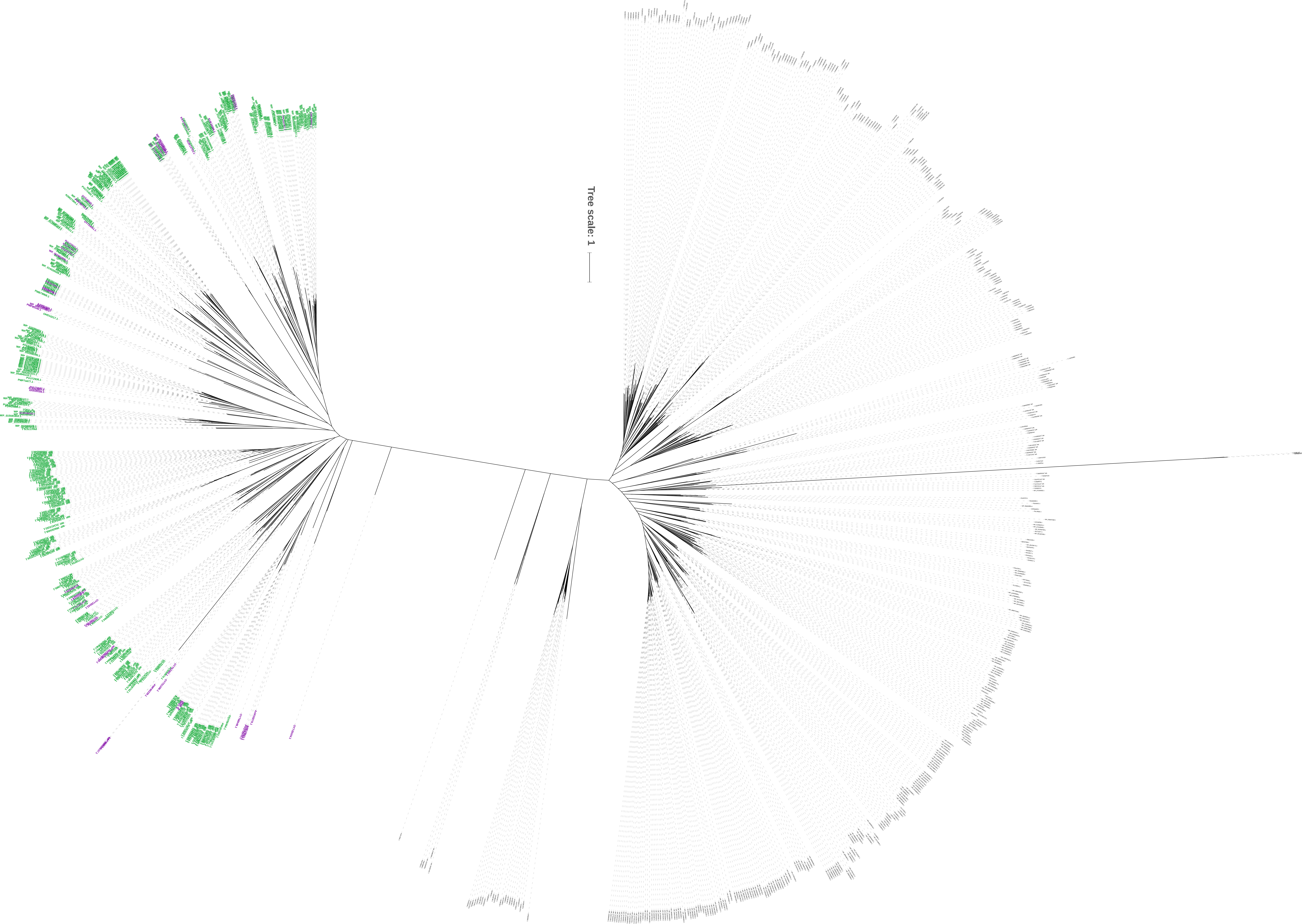
Tree scale: 1

**SI Fig. 2.** Phylogenetic tree generated from full HMMer hit results. Sequences identified as likely RG proteins by alignment with Swissprot sequences are labelled in green; sequences clustering with RG, but excluded by the alignment step are coloured in purple. Non-RG sequences (topoisomerase and helicase) sequences are labelled in black.
