## Supplementary material for "Positively twisted: The complex evolutionary history of Reverse Gyrase suggests a non-hyperthermophilic Last Universal Common Ancestor": SI Fig. 3

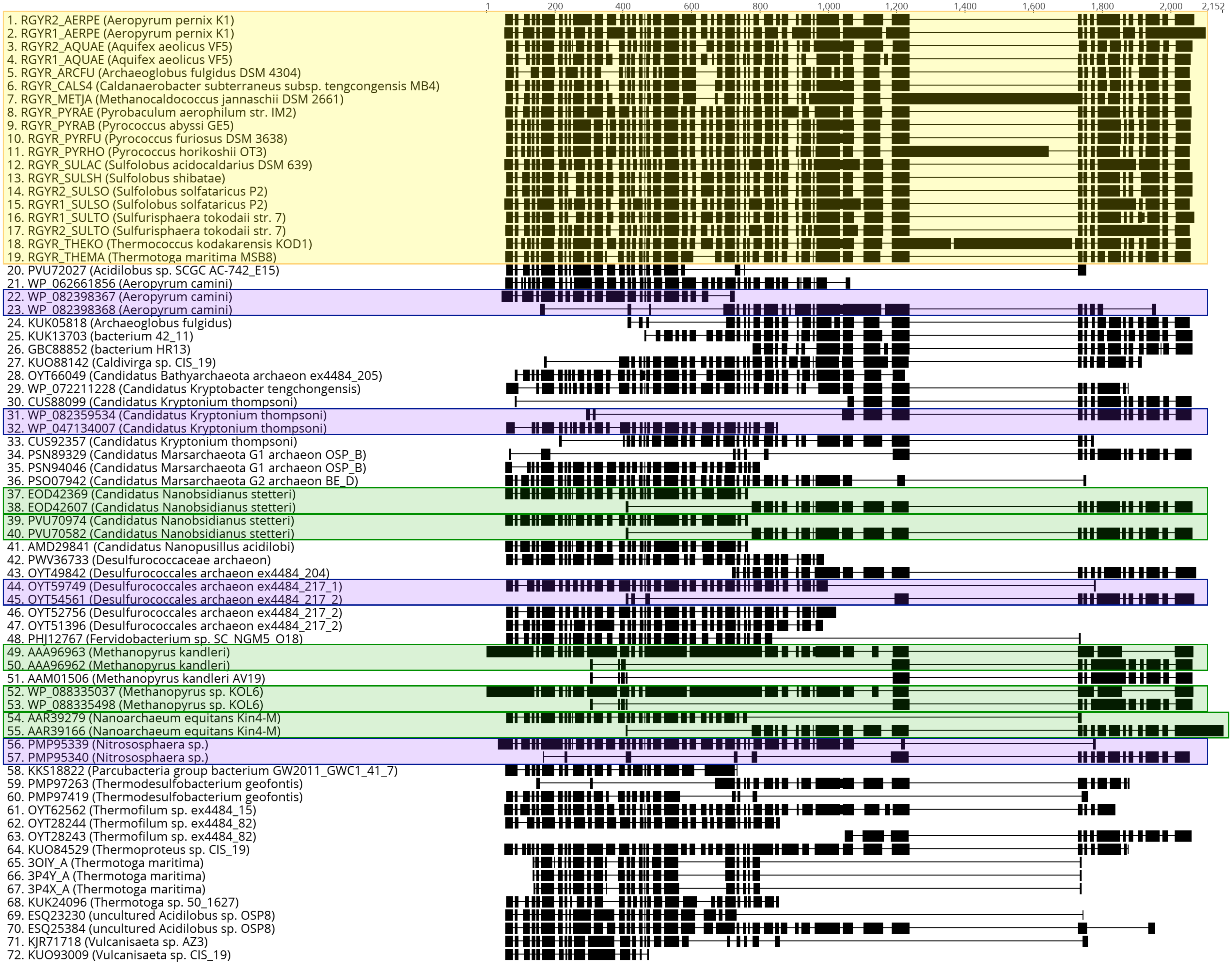
1
200
400
600
800
1,000
1,200
1,400
1,600
1,800
2,000
2,152
1. RGYR2_AERPE (Aeropyrum pernix K1)
2. RGYR1_AERPE (Aeropyrum pernix K1)
3. RGYR2_AQUAE (Aquifex aeolicus VF5)
4. RGYR1_AQUAE (Aquifex aeolicus VF5)
5. RGYR_ARCFU (Archaeoglobus fulgidus DSM 4304)
6. RGYR_CALS4 (Caldanaerobacter subterraneus subsp. tengcongensis MB4)
7. RGYR_METJA (Methanocaldococcus jannaschii DSM 2661)
8. RGYR_PYRAE (Pyrobaculum aerophilum str. IM2)
9. RGYR_PYRAB (Pyrococcus abyssi GE5)
10. RGYR_PYRFU (Pyrococcus furiosus DSM 3638)
11. RGYR_PYRHO (Pyrococcus horikoshii OT3)
12. RGYR_SULAC (Sulfolobus acidocaldarius DSM 639)
13. RGYR_SULSH (Sulfolobus shibatae)
14. RGYR2_SULSO (Sulfolobus solfataricus P2)
15. RGYR1_SULSO (Sulfolobus solfataricus P2)
16. RGYR1_SULTO (Sulfurisphaera tokodaii str. 7)
17. RGYR2_SULTO (Sulfurisphaera tokodaii str. 7)
18. RGYR_THEKO (Thermococcus kodakarensis KOD1)
19. RGYR_THEMA (Thermotoga maritima MSB8)
20. PVU72027 (Acidilobus sp. SCGC AC-742_E15)
21. WP_062661856 (Aeropyrum camini)
22. WP_082398367 (Aeropyrum camini)
23. WP_082398368 (Aeropyrum camini)
24. KUK05818 (Archaeoglobus fulgidus)
25. KUK13703 (bacterium 42_11)
26. GBC88852 (bacterium HR13)
27. KUO88142 (Caldivirga sp. CIS_19)
28. OYT66049 (Candidatus Bathyarchaeota archaeon ex4484_205)
29. WP_072211228 (Candidatus Kryptobacter tengchongensis)
30. CUS88099 (Candidatus Kryptonium thompsoni)
31. WP_082359534 (Candidatus Kryptonium thompsoni)
32. WP_047134007 (Candidatus Kryptonium thompsoni)
33. CUS92357 (Candidatus Kryptonium thompsoni)
34. PSN89329 (Candidatus Marsarchaeota G1 archaeon OSP_B)
35. PSN94046 (Candidatus Marsarchaeota G1 archaeon OSP_B)
36. PSO07942 (Candidatus Marsarchaeota G2 archaeon BE_D)
37. EOD42369 (Candidatus Nanobsidianus stetteri)
38. EOD42607 (Candidatus Nanobsidianus stetteri)
39. PVU70974 (Candidatus Nanobsidianus stetteri)
40. PVU70582 (Candidatus Nanobsidianus stetteri)
41. AMD29841 (Candidatus Nanopusillus acidilobi)
42. PWV36733 (Desulfurococcaceae archaeon)
43. OYT49842 (Desulfurococcales archaeon ex4484_204)
44. OYT59749 (Desulfurococcales archaeon ex4484_217_1)
45. OYT54561 (Desulfurococcales archaeon ex4484_217_2)
46. OYT52756 (Desulfurococcales archaeon ex4484_217_2)
47. OYT51396 (Desulfurococcales archaeon ex4484_217_2)
48. PHJ12767 (Fervidobacterium sp. SC_NGM5_O18)
49. AAA96963 (Methanopyrus kandleri)
50. AAA96962 (Methanopyrus kandleri)
51. AAM01506 (Methanopyrus kandleri AV19)
52. WP_088335037 (Methanopyrus sp. KOL6)
53. WP_088335498 (Methanopyrus sp. KOL6)
54. AAR39279 (Nanoarchaeum equitans Kin4-M)
55. AAR39166 (Nanoarchaeum equitans Kin4-M)
56. PMP95339 (Nitrososphaera sp.)
57. PMP95340 (Nitrososphaera sp.)
58. KKS18822 (Parcubacteria group bacterium GW2011_GWC1_41_7)
59. PMP97263 (Thermodesulfobacterium geofontis)
60. PMP97419 (Thermodesulfobacterium geofontis)
61. OYT62562 (Thermofilum sp. ex4484_15)
62. OYT28244 (Thermofilum sp. ex4484_82)
63. OYT28243 (Thermofilum sp. ex4484_82)
64. KUO84529 (Thermoproteus sp. CIS_19)
65. 3OIY_A (Thermotoga maritima)
66. 3P4Y_A (Thermotoga maritima)
67. 3P4X_A (Thermotoga maritima)
68. KUK24096 (Thermotoga sp. 50_1627)
69. ESQ23230 (uncultured Acidilobus sp. OSP8)
70. ESQ25384 (uncultured Acidilobus sp. OSP8)
71. KJR71718 (Vulcanisaeta sp. AZ3)
72. KUO93009 (Vulcanisaeta sp. CIS_19)

**SI Fig. 3.** Schematic of alignment of potential RG sequences identified during phylogenetic analyses which were excluded during alignment analyses of HMMer hits (see materials and methods). Protein sequence annotations and their respective species are indicated on the left, a schematic of sequence alignment shown on the right. Black boxes indicate regions with aligned residues, horizontal lines indicate gaps in alignment. Swissprot sequences are highlighted in yellow for reference of *bona fide* RG sequences. Sequences highlighted in green represent known, or anticipated split RG sequences (Nanoarchaea and *Methanopyrus* species). Sequences highlighted in purple indicate potential new split RG sequences. Non-highlighted sequences are likely truncated, or misannotated RG sequences.
