## Supplementary material for "Positively twisted: The complex evolutionary history of Reverse Gyrase suggests a non-hyperthermophilic Last Universal Common Ancestor": SI Fig. 4

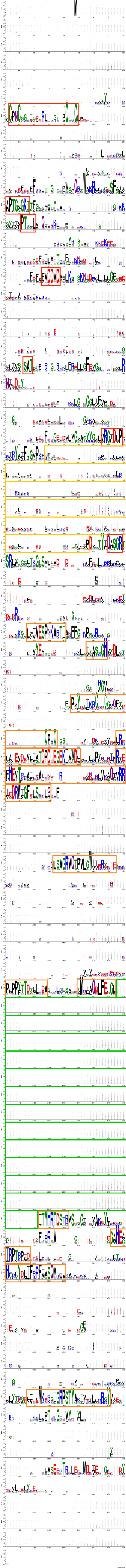
WebLogo 3.6.0

**SI Fig. 4.** Sequence logo representing the alignment of all 376 RG sequences. Regions of interest are highlighted, with conserved helicase motifs (2) in red: position 242-264 Zn finger motif; 441-448 motif I (Walker A); 486-490 motif Ia; 613-618 motif II (Walker B); 807-809 motif III; 993-999 motif V; 1235-1239 motif VI; conserved topoisomerase motifs (3) in orange: position 1408-1420 motif I; 1468-1474 motif 2; 1583-1657 Zn finger motif; 1678-1705 motif 3; 1718-1775 motif 4; 1937-1957 motif 5; 2201-2214 motif 6; 2225-2237 motif 7a; 2732-2742 motif 7b (containing the catalytic tyrosine at position 2734); 2796-2808 motif 8; 2841-2860 motif 9; 3128-3155 motif 10; poorly conserved latch domain indicated in yellow; intein indicated in green.
