## Supplementary material for "Positively twisted: The complex evolutionary history of Reverse Gyrase suggests a non-hyperthermophilic Last Universal Common Ancestor": SI Fig. 5

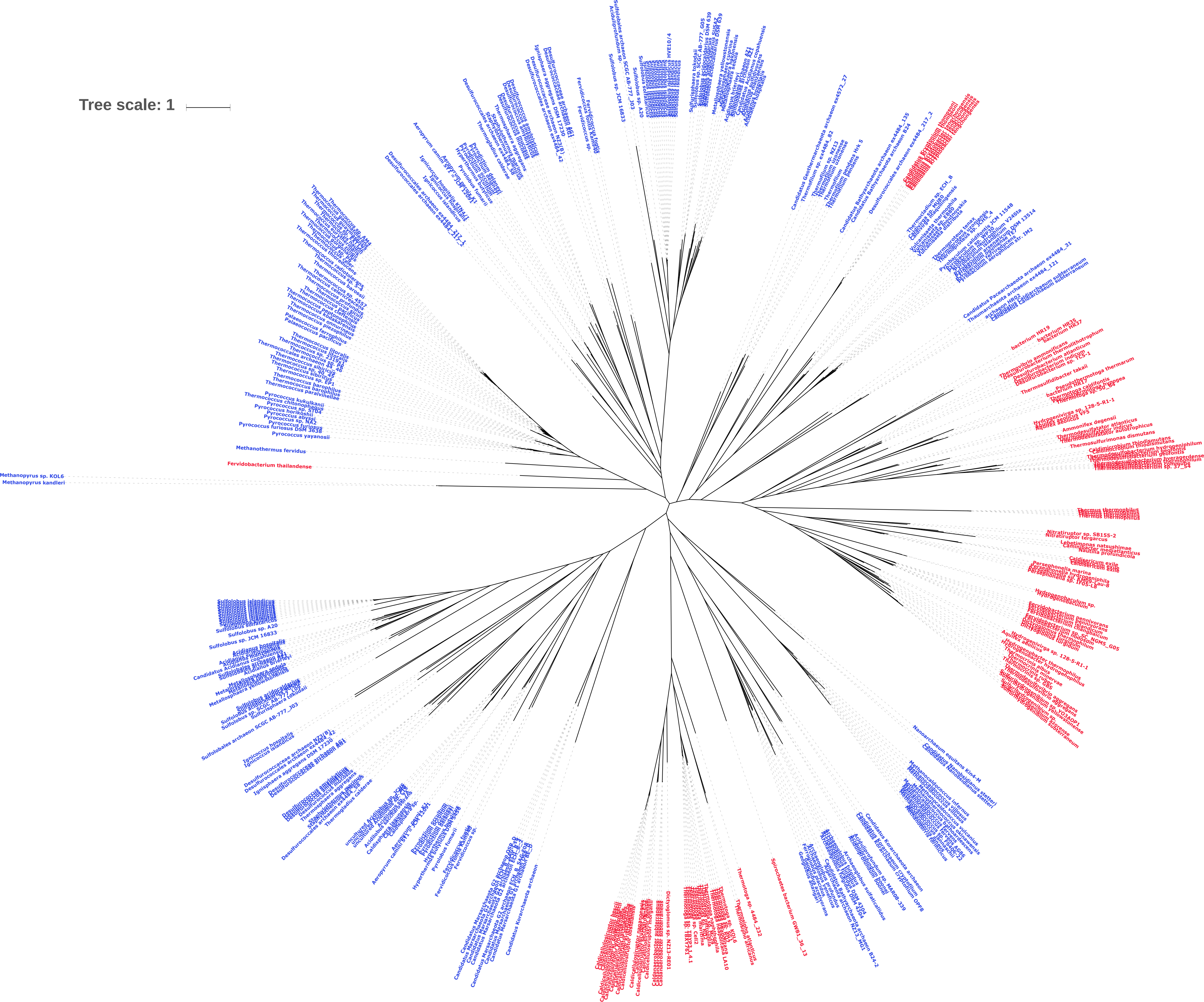
Tree scale: 1
Methanopyrus sp. KOL6
Methanopyrus kandleri
Fervidobacterium thailandense
Methanothermus fervidus
Pyrococcus yayanosii
Pyrococcus furiosus DSM 3638
Thermotoga sp. 4484_232
Spirochaetes bacterium GWB1_36_13
Dictyoglomus sp. NZ13-RE01
Nanoarchaeum equitans Kin4-M
Candidatus Korarchaeota archaeon
Persephonella marina
Hydrogenobaculum sp.
Thermosulfidibacter takaii
Pseudothermotoga thermarum
Ammonifex degensii
Thermosulfurimonas dismutans
Nitratiruptor sp. SB155-2
Sulfolobales archaeon SCGC AB-777_J03
Sulfolobus sp. JCM 16833
Sulfolobus sp. A20
Candidatus Geothermarchaeota archaeon ex4572_27
Candidatus Pacearchaeota archaeon ex4484_31

**SI Fig. 5.** Phylogenetic reconstruction of entire RG dataset using minimal trimming of sequence alignment. Trimming was performed by Noisy, removing only 17% of positions. Branch labels show species source of RG sequence, with Archaea in blue and Bacteria in red.
