## Supplementary material for "Positively twisted: The complex evolutionary history of Reverse Gyrase suggests a non-hyperthermophilic Last Universal Common Ancestor": SI Fig. 6

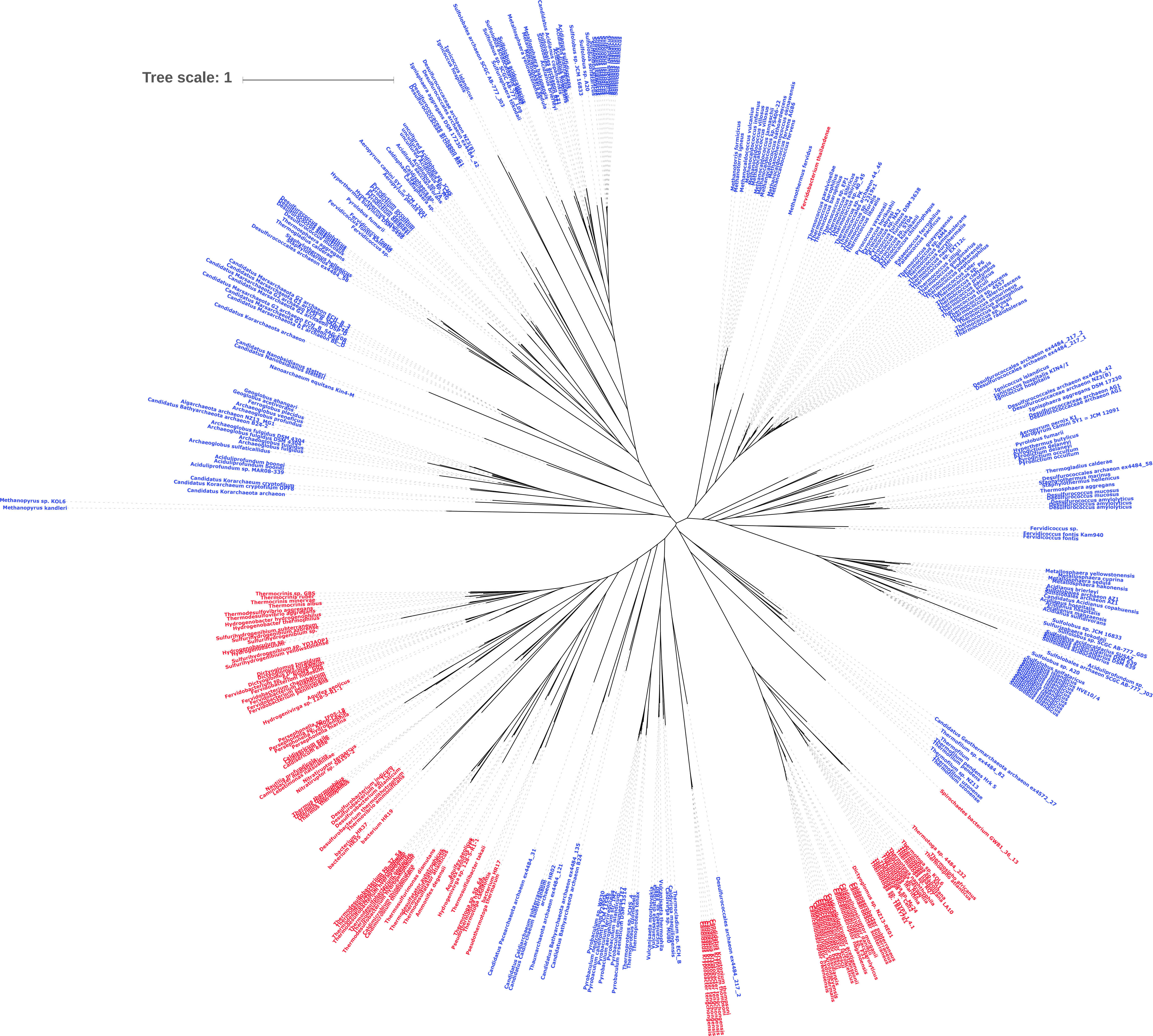
Tree scale: 1
Methanopyrus sp. KOL6
Methanopyrus kandleri
Fervidicoccus sp.
Fervidicoccus fontis Kam940
Fervidicoccus fontis
Fervidobacterium thailandense
Methanothermus fervidus
Spirochaetes bacterium GWB1_36_13

**SI Fig. 6.** Phylogenetic reconstruction of entire RG dataset using BMGE for trimming. Branch labels show species source of RG sequence, with Archaea in blue and Bacteria in red.
