## Supplementary material for "Positively twisted: The complex evolutionary history of Reverse Gyrase suggests a non-hyperthermophilic Last Universal Common Ancestor": SI Fig. 7

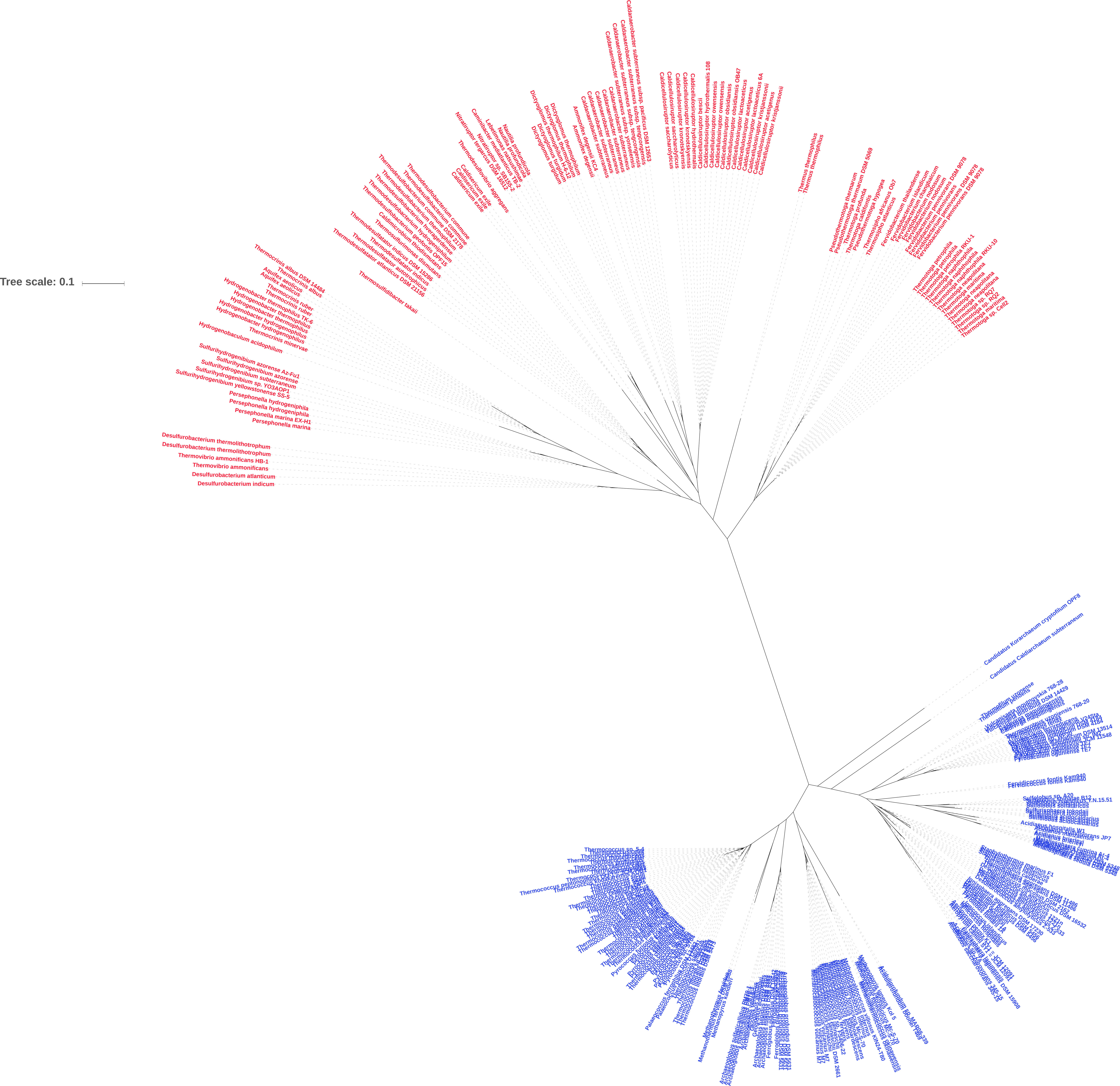

Tree scale: 0.1
Thermocrinis albus DSM 14484
Thermocrinis albus
Aquifex aeolicus
Thermocrinis ruber
Hydrogenobacter thermophilus TK-6
Hydrogenobacter thermophilus
Hydrogenobacter hydrogenophilus
Thermocrinis minervae
Hydrogenobaculum acidophilum
Sulfurihydrogenibium azorense Az-Fu1
Sulfurihydrogenibium azorense
Sulfurihydrogenibium subterraneum
Sulfurihydrogenibium sp. YO3AOP1
Sulfurihydrogenibium yellowstonense SS-5
Persephonella hydrogeniphila
Persephonella marina EX-H1
Persephonella marina
Desulfurobacterium thermolithotrophum
Thermovibrio ammonificans HB-1
Thermovibrio ammonificans
Desulfurobacterium atlanticum
Desulfurobacterium indicum
Thermosulfidibacter takaii
Thermodesulfatator atlanticus DSM 21156
Thermodesulfatator autotrophicus
Thermodesulfatator indicus
Thermodesulfatator indicus DSM 15286
Thermosulfurimonas dismutans
Caldimicrobium thiodismutans
Thermodesulfobacterium geofontis OPF15
Thermodesulfobacterium hydrogeniphilum
Thermodesulfobacterium hveragerdense
Thermodesulfobacterium commune DSM 2178
Thermodesulfobacterium commune
Caldisericum exile
Thermodesulfovibrio aggregans
Nitratiruptor tergarcus DSM 16512
Nitratiruptor sp. SB155-2
Caminibacter mediatlanticus TB-2
Lebetimonas natsushimae
Nautilia profundicola
Dictyoglomus turgidum
Dictyoglomus thermophilum H-6-12
Dictyoglomus thermophilum
Ammonifex degensii
Ammonifex degensii KC4
Caldanaerobacter subterraneus
Caldanaerobacter subterraneus subsp. yonseiensis
Caldanaerobacter subterraneus subsp. tengcongensis
Caldanaerobacter subterraneus subsp. pacificus DSM 12653
Caldicellulosiruptor saccharolyticus
Caldicellulosiruptor kronotskyensis
Caldicellulosiruptor hydrothermalis
Caldicellulosiruptor hydrothermalis 108
Caldicellulosiruptor bescii
Caldicellulosiruptor owensensis
Caldicellulosiruptor obsidiansis
Caldicellulosiruptor obsidiansis OB47
Caldicellulosiruptor lactoaceticus
Caldicellulosiruptor acetigenus
Caldicellulosiruptor lactoaceticus 6A
Caldicellulosiruptor kristjanssonii
Thermus thermophilus
Pseudothermotoga thermarum
Pseudothermotoga thermarum DSM 5069
Thermotoga profunda
Thermotoga caldifontis
Pseudothermotoga hypogea
Thermosipho africanus Ob7
Thermosipho atlanticus
Fervidobacterium thailandense
Fervidobacterium islandicum
Fervidobacterium changbaicum
Fervidobacterium nodosum
Fervidobacterium pennivorans
Fervidobacterium pennivorans DSM 9078
Thermotoga petrophila
Thermotoga petrophila RKU-1
Thermotoga naphthophila
Thermotoga naphthophila RKU-10
Thermotoga maritima
Thermotoga neapolitana
Thermotoga sp. RQ2
Thermotoga sp. Cell2
Candidatus Korarchaeum cryptofilum OPF8
Candidatus Caldiarchaeum subterraneum
Fervidicoccus fontis Kam940
Sulfolobus solfataricus
Acidianus hospitalis W1
Staphylothermus marinus F1
Thermococcus sp. 4557
Palaeococcus pacificus
Methanopyrus kandleri
Archaeoglobus profundus DSM 5631
Acidilobus saccharovorans 345-15

**SI Fig. 7.** Phylogenetic reconstruction of 16S rDNA gene from RG-encoding species. Sequences encoded by Archaeal species are indicated in blue, Bacteria in red.
