## Supplementary material for "Positively twisted: The complex evolutionary history of Reverse Gyrase suggests a non-hyperthermophilic Last Universal Common Ancestor": SI Fig. 8

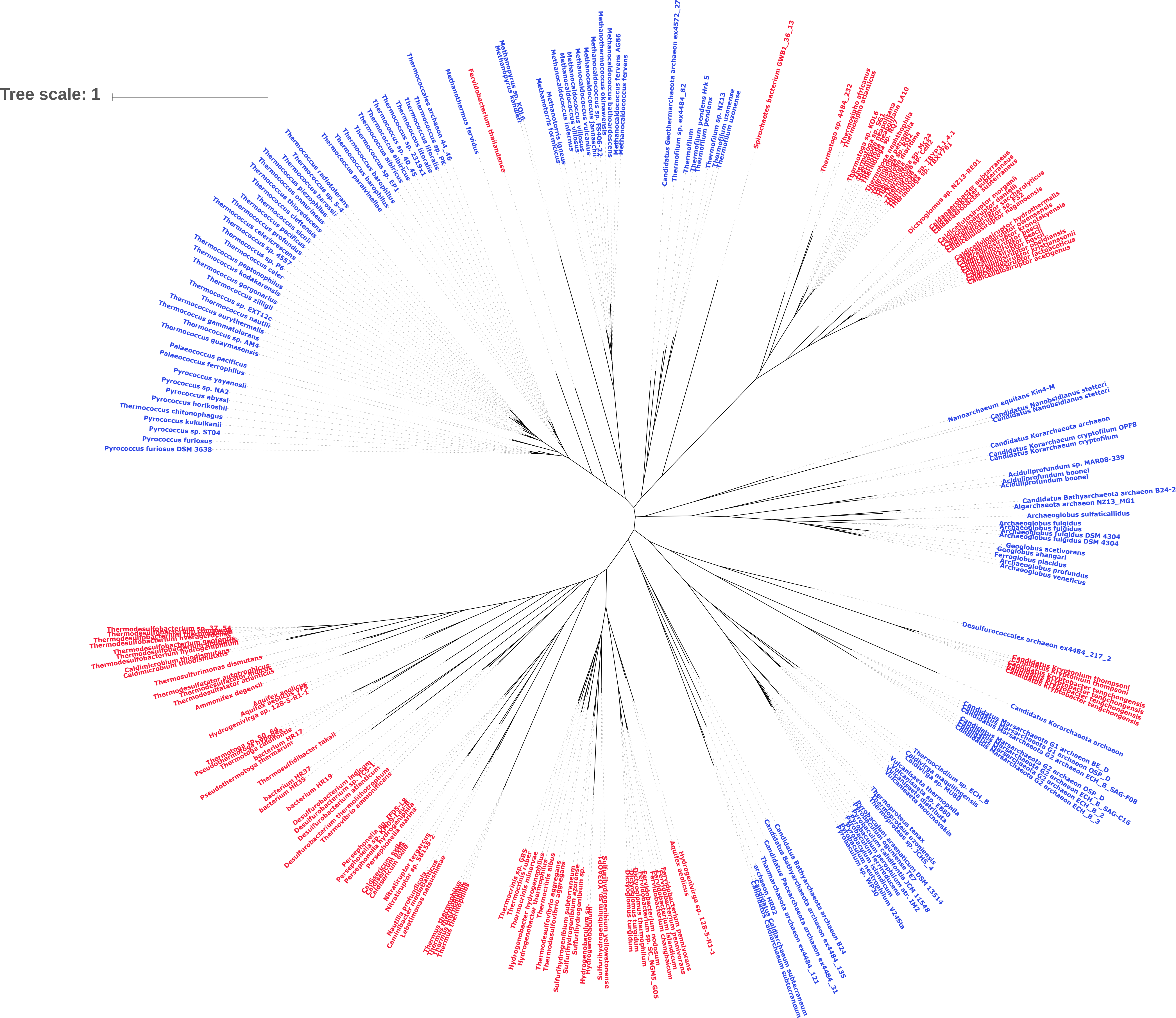

Tree scale: 1
Pyrococcus furiosus DSM 3638
Pyrococcus furiosus
Pyrococcus sp. ST04
Pyrococcus kukulkanii
Thermococcus chitonophagus
Pyrococcus horikoshii
Pyrococcus abyssi
Pyrococcus sp. NA2
Pyrococcus yayanosii
Palaeococcus ferrophilus
Palaeococcus pacificus
Thermococcus guaymasensis
Thermococcus sp. AM4
Thermococcus gammatolerans
Thermococcus eurythermalis
Thermococcus nautili
Thermococcus sp. EXT12c
Thermococcus zilligii
Thermococcus gorgonarius
Thermococcus kodakarensis
Thermococcus peptonophilus
Thermococcus celer
Thermococcus sp. P6
Thermococcus sp. 4557
Thermococcus celericrescens
Thermococcus profundus
Thermococcus pacificus
Thermococcus siculi
Thermococcus cleftensis
Thermococcus onnurineus
Thermococcus thioreducens
Thermococcus piezophilus
Thermococcus barossii
Thermococcus sp. 5-4
Thermococcus radiotolerans
Thermococcus paralvinellae
Thermococcus barophilus
Thermococcus sp. EP1
Thermococcus sibiricus
Thermococcus sp. 40_45
Thermococcus sp. 2319x1
Thermococcus litoralis
Thermococcus sp. PK
Thermococcales archaeon 44_46
Methanothermus fervidus
Fervidobacterium thailandense
Methanopyrus kandleri
Methanopyrus sp. KOL6
Methanotorris formicicus
Methanotorris igneus
Methanocaldococcus infernus
Methanocaldococcus villosus
Methanocaldococcus vulcanius
Methanocaldococcus jannaschii
Methanocaldococcus sp. FS406-22
Methanothermococcus okinawensis
Methanocaldococcus bathoardescens
Methanocaldococcus fervens AG86
Methanocaldococcus fervens
Candidatus Geothermarchaeota archaeon ex4572_27
Thermofilum sp. ex4484_82
Thermofilum
Thermofilum pendens Hrk 5
Thermofilum pendens
Thermofilum sp. NZ13
Thermofilum uzonense
Thermofilum uzonense
Spirochaetes bacterium GWB1_36_13
Thermotoga sp. 4484_232
Thermosipho africanus
Thermosipho atlanticus
Thermotoga sp. KOL6
Thermotoga neapolitana LA10
Thermotoga naphthophila
Thermotoga sp. Mc24
Thermotoga sp. TBYP3.1.4.1
Thermotoga sp. Cell2
Thermotoga sp. RQ7
Thermotoga sp. 1BA7761
Dictyoglomus sp. NZ13-RE01
Caldanaerobacter subterraneus
Caldanaerobacter subterraneus
Caldicellulosiruptor morganii
Caldicellulosiruptor danielii
Caldicellulosiruptor sp. F32
Caldicellulosiruptor saccharolyticus
Caldicellulosiruptor owensensis
Caldicellulosiruptor naganoensis
Caldicellulosiruptor hydrothermalis
Caldicellulosiruptor kronotskyensis
Caldicellulosiruptor bescii
Caldicellulosiruptor obsidiansis
Caldicellulosiruptor kristjanssonii
Caldicellulosiruptor lactoaceticus
Caldicellulosiruptor acetigenus
Nanoarchaeum equitans Kin4-M
Candidatus Nanobsidianus stetteri
Candidatus Nanobsidianus stetteri
Candidatus Korarchaeota archaeon
Candidatus Korarchaeum cryptofilum OPF8
Candidatus Korarchaeum cryptofilum
Aciduliprofundum sp. MAR08-339
Aciduliprofundum boonei
Aciduliprofundum boonei
Candidatus Bathyarchaeota archaeon B24-2
Aigarchaeota archaeon NZ13_MG1
Archaeoglobus sulfaticallidus
Archaeoglobus fulgidus
Archaeoglobus fulgidus
Archaeoglobus fulgidus DSM 4304
Archaeoglobus fulgidus DSM 4304
Geoglobus acetivorans
Geoglobus ahangari
Ferroglobus placidus
Archaeoglobus profundus
Archaeoglobus veneficus
Desulfurococcales archaeon ex4484_217_2
Candidatus Kryptonium thompsoni
Candidatus Kryptonium thompsoni
Candidatus Kryptobacter tengchongensis
Candidatus Kryptobacter tengchongensis
Candidatus Kryptobacter tengchongensis
Candidatus Korarchaeota archaeon
Candidatus Marsarchaeota G1 archaeon BE_D
Candidatus Marsarchaeota G1 archaeon OSP_D
Candidatus Marsarchaeota G1 archaeon ECH_B_SAG-F08
Candidatus Marsarchaeota G2 archaeon OSP_D
Candidatus Marsarchaeota G2 archaeon ECH_B_SAG-C16
Candidatus Marsarchaeota G2 archaeon ECH_B_2
Candidatus Marsarchaeota G2 archaeon ECH_B_3
Thermocladium sp. ECH_B
Caldivirga maquilingensis
Caldivirga sp. MU80
Vulcanisaeta thermophila
Vulcanisaeta sp. EB80
Vulcanisaeta distributa
Vulcanisaeta moutnovskia
Thermoproteus tenax
Thermoproteus uzoniensis
Thermoproteus sp. JCHS_4
Pyrobaculum arsenaticum DSM 13514
Pyrobaculum oguniense TE7
Pyrobaculum calidifontis JCM 11548
Pyrobaculum aerophilum str. IM2
Pyrobaculum ferrireducens
Pyrobaculum neutrophilum V24Sta
Pyrobaculum sp. WP30
Candidatus Bathyarchaeota archaeon B24
Candidatus Bathyarchaeota archaeon ex4484_135
Candidatus Pacearchaeota archaeon ex4484_31
Thaumarchaeota archaeon ex4484_121
archaeon HR02
Candidatus Caldiarchaeum subterraneum
Candidatus Caldiarchaeum subterraneum
Hydrogenivirga sp. 128-5-R1-1
Aquifex aeolicus
Fervidobacterium pennivorans
Fervidobacterium pennivorans
Fervidobacterium islandicum
Fervidobacterium changbaicum
Fervidobacterium nodosum
Fervidobacterium sp. SC_NGMS_G05
Dictyoglomus thermophilum
Dictyoglomus turgidum
Dictyoglomus turgidum
Sulfurihydrogenibium yellowstonense
Sulfurihydrogenibium sp. YO3AOP1
Hydrogenobaculum sp.
Hydrogenobaculum sp.
Sulfurihydrogenibium azorense
Sulfurihydrogenibium subterraneum
Thermodesulfovibrio aggregans
Thermodesulfovibrio aggregans
Thermocrinis albus
Hydrogenobacter thermophilus
Hydrogenobacter hydrogenophilus
Thermocrinis minervae
Thermocrinis ruber
Thermocrinis sp. GBS
Thermus thermophilus
Lebetimonas natsushimae
Caminibacter mediatlanticus
Nautilia profundicola
Nitratiruptor sp. SB155-2
Nitratiruptor tergarcus
Caldisericum exile
Caldisericum exile
Persephonella marina
Persephonella hydrogeniphila
Persephonella sp. KM09-Lau-8
Persephonella sp. IF05-L8
Desulfurobacterium thermolithotrophum
Thermovibrio ammonificans
Desulfurobacterium atlanticum
Desulfurobacterium sp. TC5-1
Desulfurobacterium indicum
bacterium HR19
bacterium HR35
bacterium HR37
Thermosulfidibacter takaii
Pseudothermotoga thermarum
bacterium HR17
Pseudothermotoga hypogea
Thermotoga caldifontis
Thermotoga sp. 50_64
Hydrogenivirga sp. 128-5-R1-1
Aquifex aeolicus VF5
Aquifex aeolicus
Ammonifex degensii
Thermodesulfatator atlanticus
Thermodesulfatator indicus
Thermodesulfatator autotrophicus
Thermosulfurimonas dismutans
Caldimicrobium thiodismutans
Caldimicrobium thiodismutans
Thermodesulfobacterium hydrogeniphilum
Thermodesulfobacterium geofontis
Thermodesulfobacterium hveragerdense
Thermodesulfobacterium commune
Thermodesulfobacterium sp. 37_54

**SI Fig. 8.** Phylogenetic reconstruction of RG dataset with TopR1-like and TopR2-like sequences removed. Branch labels show species source of RG sequence, with Archaea in blue and Bacteria in red.
