## Supplementary material for "Positively twisted: The complex evolutionary history of Reverse Gyrase suggests a non-hyperthermophilic Last Universal Common Ancestor": SI Fig. 9

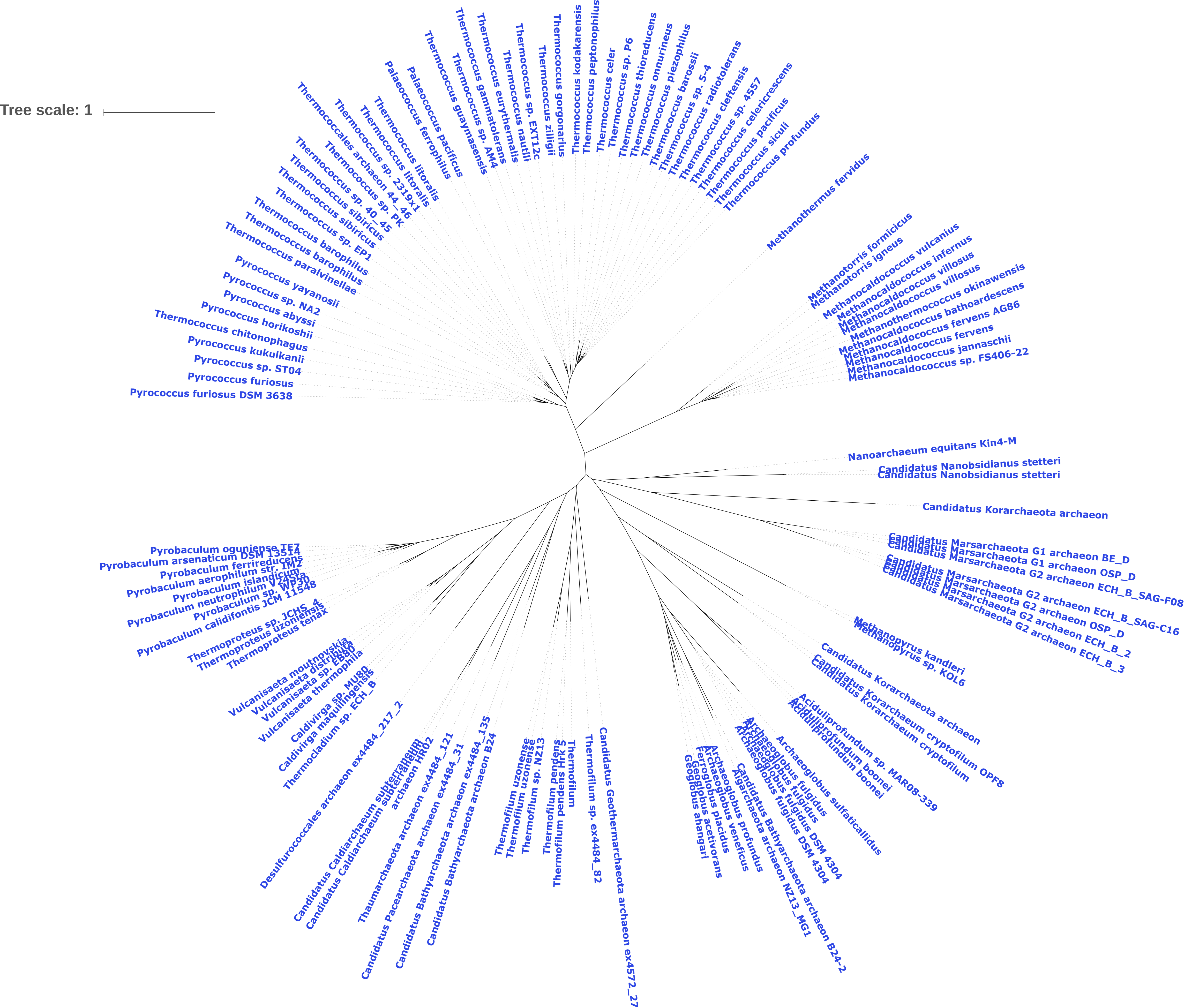
Tree scale: 1
Pyrococcus furiosus DSM 3638
Pyrococcus furiosus
Pyrococcus sp. ST04
Pyrococcus kukulkanii
Thermococcus chitonophagus
Pyrococcus horikoshii
Pyrococcus abyssi
Pyrococcus sp. NA2
Pyrococcus yayanosii
Thermococcus paralvinellae
Thermococcus barophilus
Thermococcus barophilus
Thermococcus sp. EP1
Thermococcus sibiricus
Thermococcus sibiricus
Thermococcus sp. 40_45
Thermococcus sp. PK
Thermococcales archaeon 44_46
Thermococcus sp. 2319x1
Thermococcus litoralis
Thermococcus litoralis
Palaeococcus ferrophilus
Palaeococcus pacificus
Thermococcus guaymasensis
Thermococcus sp. AM4
Thermococcus gammatolerans
Thermococcus eurythermalis
Thermococcus nautili
Thermococcus sp. EXT12c
Thermococcus zilligii
Thermococcus gorgonarius
Thermococcus kodakarensis
Thermococcus peptonophilus
Thermococcus celer
Thermococcus sp. P6
Thermococcus thioreducens
Thermococcus onnurineus
Thermococcus piezophilus
Thermococcus barossii
Thermococcus sp. 5-4
Thermococcus radiotolerans
Thermococcus cleftensis
Thermococcus sp. 4557
Thermococcus celericrescens
Thermococcus pacificus
Thermococcus siculi
Thermococcus profundus
Methanothermus fervidus
Methanotorris formicicus
Methanotorris igneus
Methanocaldococcus vulcanius
Methanocaldococcus infernus
Methanocaldococcus villosus
Methanocaldococcus villosus
Methanothermococcus okinawensis
Methanocaldococcus bathoardescens
Methanocaldococcus fervens AG86
Methanocaldococcus fervens
Methanocaldococcus jannaschii
Methanocaldococcus sp. FS406-22
Nanoarchaeum equitans Kin4-M
Candidatus Nanobsidianus stetteri
Candidatus Nanobsidianus stetteri
Candidatus Korarchaeota archaeon
Candidatus Marsarchaeota G1 archaeon BE_D
Candidatus Marsarchaeota G1 archaeon OSP_D
Candidatus Marsarchaeota G2 archaeon ECH_B_SAG-F08
Candidatus Marsarchaeota G2 archaeon ECH_B_SAG-C16
Candidatus Marsarchaeota G2 archaeon OSP_D
Candidatus Marsarchaeota G2 archaeon ECH_B_2
Candidatus Marsarchaeota G2 archaeon ECH_B_3
Methanopyrus kandleri
Methanopyrus sp. KOL6
Candidatus Korarchaeota archaeon
Candidatus Korarchaeum cryptofilum OPF8
Candidatus Korarchaeum cryptofilum
Aciduliprofundum sp. MAR08-339
Aciduliprofundum boonei
Aciduliprofundum boonei
Archaeoglobus sulfaticallidus
Archaeoglobus fulgidus
Archaeoglobus fulgidus DSM 4304
Archaeoglobus fulgidus DSM 4304
Candidatus Bathyarchaeota archaeon B24-2
Aigarchaeota archaeon NZ13_MG1
Archaeoglobus profundus
Archaeoglobus veneficus
Geoglobus acetivorans
Ferroglobus placidus
Geoglobus ahangari
Candidatus Geothermarchaeota archaeon ex4572_27
Thermofilum sp. ex4484_82
Thermofilum
Thermofilum pendens Hrk 5
Thermofilum pendens
Thermofilum sp. NZ13
Thermofilum uzonense
Thermofilum uzonense
Candidatus Bathyarchaeota archaeon B24
Candidatus Bathyarchaeota archaeon ex4484_135
Candidatus Pacearchaeota archaeon ex4484_31
Thaumarchaeota archaeon ex4484_121
archaeon HR02
Candidatus Caldiarchaeum subterraneum
Candidatus Caldiarchaeum subterraneum
Desulfurococcales archaeon ex4484_217_2
Thermocladium sp. ECH_B
Caldivirga maquilingensis
Caldivirga sp. MU80
Vulcanisaeta thermophila
Vulcanisaeta sp. EB80
Vulcanisaeta distributa
Vulcanisaeta moutnovskia
Thermoproteus tenax
Thermoproteus uzoniensis
Thermoproteus sp. JCHS_4
Pyrobaculum calidifontis JCM 11548
Pyrobaculum sp. WP30
Pyrobaculum neutrophilum V24Sta
Pyrobaculum islandicum
Pyrobaculum aerophilum str. IM2
Pyrobaculum ferrireducens
Pyrobaculum arsenaticum DSM 13514
Pyrobaculum oguniense TE7

**SI FIg. 9.** Phylogenetic reconstruction of Archaeal RG dataset with TopR1-like and TopR2-like sequences removed. Branch labels show species source of RG sequence
