## Supplementary material for "Positively twisted: The complex evolutionary history of Reverse Gyrase suggests a non-hyperthermophilic Last Universal Common Ancestor": SI Fig. 10

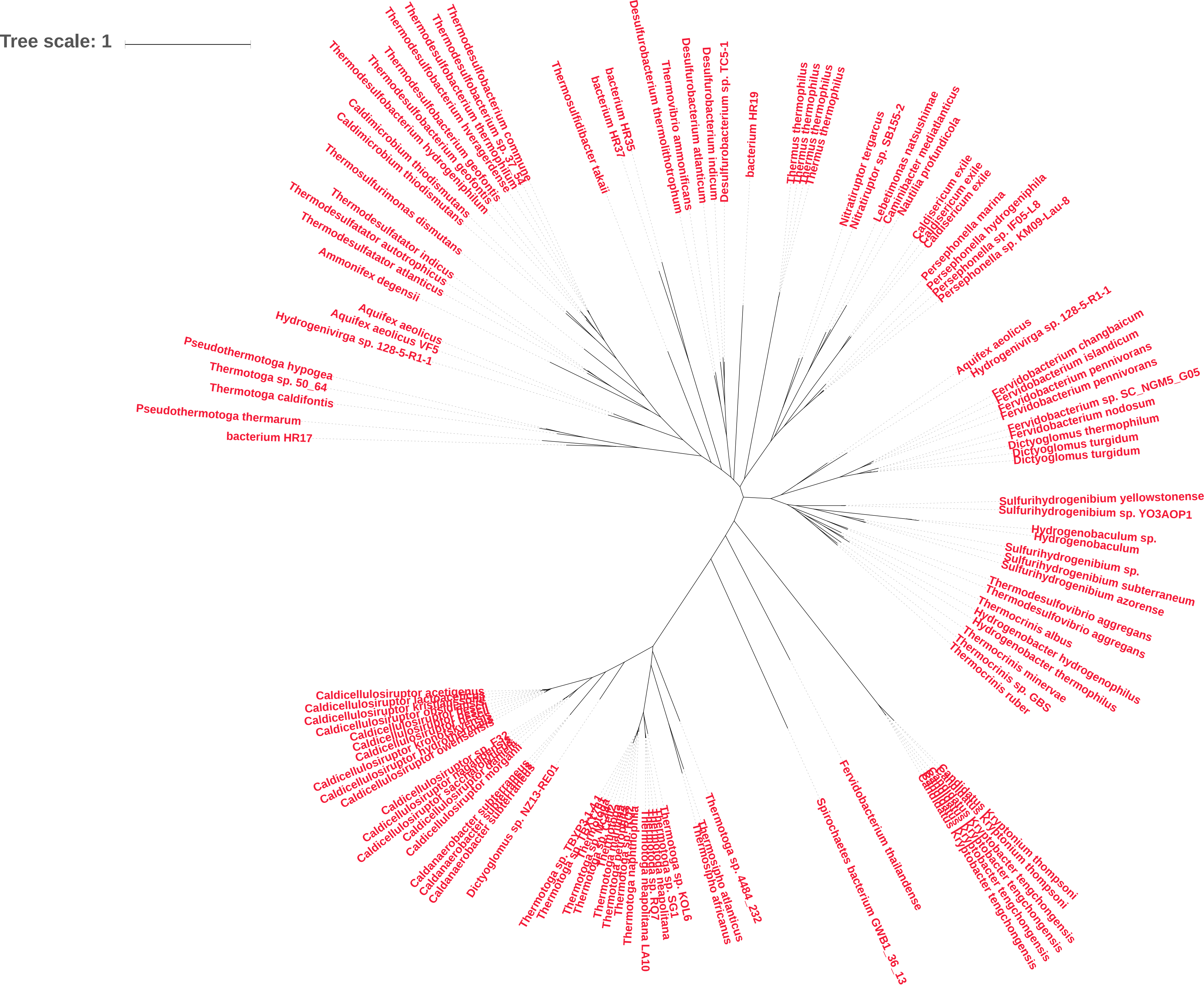

Tree scale: 1
Thermodesulfobacterium commune
Thermodesulfobacterium sp. 3_64
Thermodesulfobacterium thermophilum
Thermodesulfobacterium hveragerdense
Thermodesulfobacterium geofontis
Thermodesulfobacterium hydrogeniphilum
Caldimicrobium thiodismutans
Caldimicrobium thiodismutans
Thermosulfurimonas dismutans
Thermodesulfatator indicus
Thermodesulfatator autotrophicus
Thermodesulfatator atlanticus
Ammonifex degensii
Aquifex aeolicus
Aquifex aeolicus VF5
Hydrogenivirga sp. 128-5-R1-1
Pseudothermotoga hypogea
Thermotoga sp. 50_64
Thermotoga caldifontis
Pseudothermotoga thermarum
bacterium HR17
Thermosulfidibacter takaii
bacterium HR35
bacterium HR37
Desulfurobacterium thermolithotrophum
Thermovibrio ammonificans
Desulfurobacterium atlanticum
Desulfurobacterium indicum
Desulfurobacterium sp. TC5-1
bacterium HR19
Thermus thermophilus
Thermus thermophilus
Thermus thermophilus
Thermus thermophilus
Nitratiruptor tergarcus
Nitratiruptor sp. SB155-2
Lebetimonas natsushimae
Caminibacter mediatlanticus
Nautilia profundicola
Caldisericum exile
Caldisericum exile
Caldisericum exile
Persephonella marina
Persephonella hydrogeniphila
Persephonella sp. IF05-L8
Persephonella sp. KM09-Lau-8
Aquifex aeolicus
Hydrogenivirga sp. 128-5-R1-1
Fervidobacterium changbaicum
Fervidobacterium islandicum
Fervidobacterium pennivorans
Fervidobacterium pennivorans
Fervidobacterium sp. SC_NGM5_G05
Fervidobacterium nodosum
Dictyoglomus thermophilum
Dictyoglomus turgidum
Dictyoglomus turgidum
Sulfurihydrogenibium yellowstonense
Sulfurihydrogenibium sp. YO3AOP1
Hydrogenobaculum sp.
Hydrogenobaculum
Sulfurihydrogenibium sp.
Sulfurihydrogenibium subterraneum
Sulfurihydrogenibium azorense
Thermodesulfovibrio aggregans
Thermodesulfovibrio aggregans
Thermocrinis albus
Hydrogenobacter hydrogenophilus
Hydrogenobacter thermophilus
Thermocrinis minervae
Thermocrinis sp. GBS
Thermocrinis ruber
Candidatus Kryptonium thompsoni
Candidatus Kryptonium thompsoni
Candidatus Kryptobacter tengchongensis
Candidatus Kryptobacter tengchongensis
Candidatus Kryptobacter tengchongensis
Fervidobacterium thailandense
Spirochaetes bacterium GWB1_36_13
Thermotoga sp. 4484_232
Thermosipho atlanticus
Thermosipho africanus
Thermotoga sp. KOL6
Thermotoga sp. SG1
Thermotoga neapolitana
Thermotoga sp. RQ7
Thermotoga neapolitana LA10
Thermotoga naphthophila
Dictyoglomus sp. NZ13-RE01
Caldanaerobacter subterraneus
Caldanaerobacter subterraneus
Caldanaerobacter subterraneus
Caldicellulosiruptor morganii
Caldicellulosiruptor danielii
Caldicellulosiruptor saccharolyticus
Caldicellulosiruptor naganoensis
Caldicellulosiruptor sp. F32
Caldicellulosiruptor owensensis
Caldicellulosiruptor hydrothermalis
Caldicellulosiruptor kronotskyensis
Caldicellulosiruptor acetigenus
Caldicellulosiruptor lactoaceticus
Caldicellulosiruptor kristjanssonii
Caldicellulosiruptor obsidiansis

**SI Fig. 10.** Phylogenetic reconstruction of Bacterial RG dataset. Branch labels show species source of RG sequence
