## Supplementary material for "Positively twisted: The complex evolutionary history of Reverse Gyrase suggests a non-hyperthermophilic Last Universal Common Ancestor": SI Fig. 11

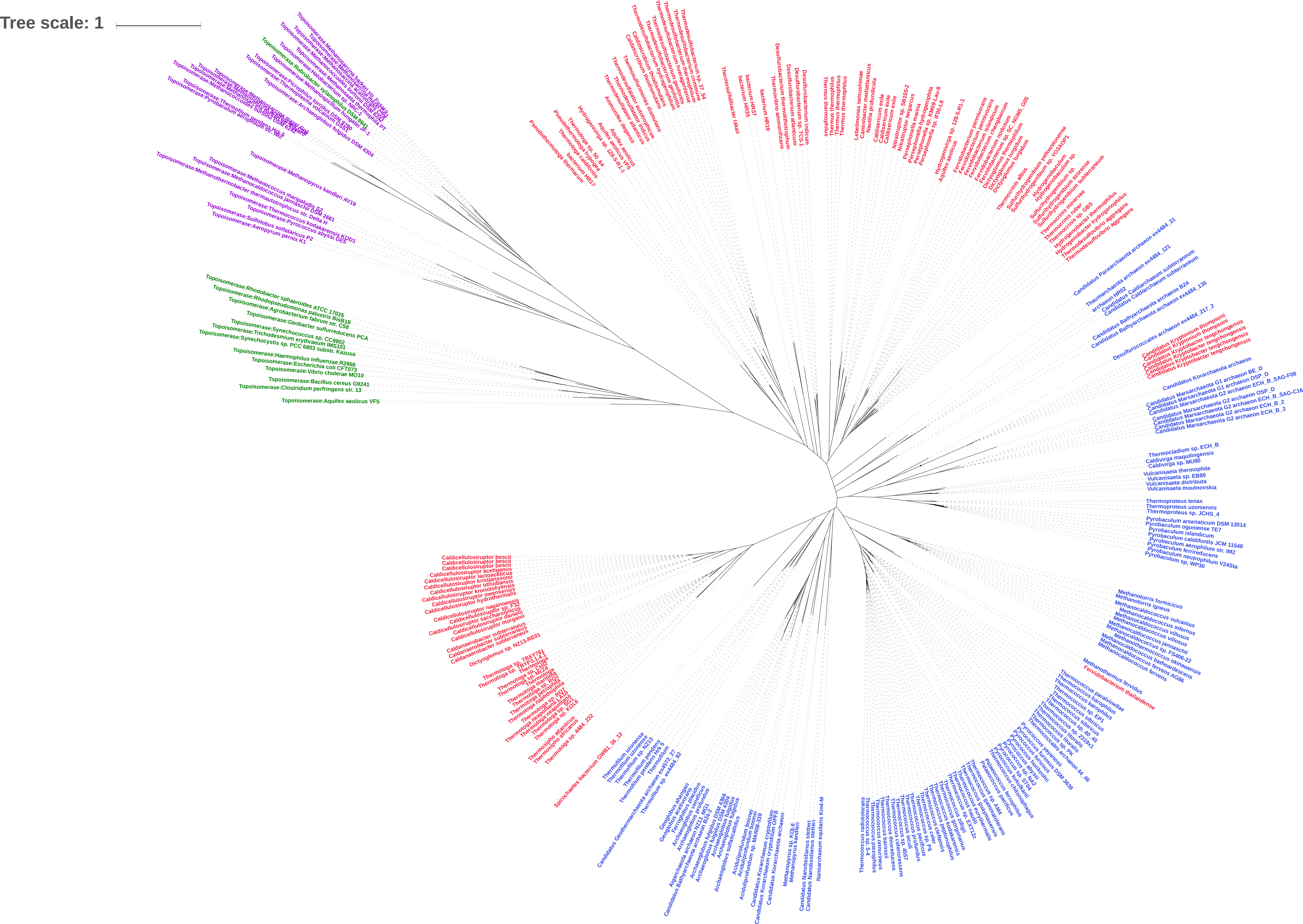
Tree scale: 1
Topoisomerase:Methanopyrus kandleri AV19
Topoisomerase:Methanococcus maripaludis S2
Topoisomerase:Methanocaldococcus jannaschii DSM 2661
Topoisomerase:Methanothermobacter thermautotrophicus str. Delta H
Topoisomerase:Thermococcus kodakarensis KOD1
Topoisomerase:Pyrococcus abyssi GE5
Topoisomerase:Sulfolobus solfataricus P2
Topoisomerase:Aeropyrum pernix K1
Topoisomerase:Rhodobacter sphaeroides ATCC 17025
Topoisomerase:Rhodopseudomonas palustris BisB18
Topoisomerase:Agrobacterium fabrum str. C58
Topoisomerase:Geobacter sulfurreducens PCA
Topoisomerase:Synechococcus sp. CC9902
Topoisomerase:Trichodesmium erythraeum IMS101
Topoisomerase:Synechocystis sp. PCC 6803 substr. Kazusa
Topoisomerase:Haemophilus influenzae R2866
Topoisomerase:Escherichia coli CFT073
Topoisomerase:Vibrio cholerae MO10
Topoisomerase:Bacillus cereus G9241
Topoisomerase:Clostridium perfringens str. 13
Topoisomerase:Aquifex aeolicus VF5
Candidatus Pacearchaeota archaeon ex4484_31
Thaumarchaeota archaeon ex4484_121
archaeon HR02
Candidatus Caldiarchaeum subterraneum
Candidatus Caldiarchaeum subterraneum
Candidatus Bathyarchaeota archaeon B24
Candidatus Bathyarchaeota archaeon ex4484_135
Desulfurococcales archaeon ex4484_217_2
Candidatus Korarchaeota archaeon
Thermocladium sp. ECH_B
Caldivirga maquilingensis
Caldivirga sp. MU80
Vulcanisaeta thermophila
Vulcanisaeta sp. EB80
Vulcanisaeta distributa
Vulcanisaeta moutnovskia
Thermoproteus tenax
Thermoproteus uzoniensis
Thermoproteus sp. JCHS_4
Pyrobaculum arsenaticum DSM 13514
Pyrobaculum oguniense TE7
Pyrobaculum islandicum
Pyrobaculum calidifontis JCM 11548
Pyrobaculum aerophilum str. IM2
Pyrobaculum ferrireducens
Pyrobaculum neutrophilum V24Sta
Pyrobaculum sp. WP30
Methanotorris formicicus
Methanotorris igneus
Methanocaldococcus vulcanius
Methanocaldococcus infernus
Methanocaldococcus villosus
Methanocaldococcus jannaschii
Methanocaldococcus sp. FS406-22
Methanothermococcus okinawensis
Methanocaldococcus bathoardescens
Methanocaldococcus fervens AG86
Methanothermus fervidus
Fervidobacterium thailandense
Caldicellulosiruptor bescii
Caldicellulosiruptor bescii
Caldicellulosiruptor bescii
Caldicellulosiruptor acetigenus
Caldicellulosiruptor lactoaceticus
Caldicellulosiruptor kristjanssonii
Caldicellulosiruptor obsidiansis
Caldicellulosiruptor kronotskyensis
Caldicellulosiruptor owensensis
Caldicellulosiruptor hydrothermalis
Caldicellulosiruptor naganoensis
Caldicellulosiruptor sp. F32
Caldicellulosiruptor saccharolyticus
Caldicellulosiruptor danielii
Caldicellulosiruptor morganii
Caldanaerobacter subterraneus
Caldanaerobacter subterraneus
Caldanaerobacter subterraneus
Dictyoglomus sp. NZ13-RE01
Thermus thermophilus
Nanoarchaeum equitans Kin4-M

**SI Fig. 11.** Phylogenetic reconstruction of RG dataset with TopR1-like and TopR2-like sequences removed. Bacterial and Archaeal topoisomerase sequences included as an outgroup. Branch labels show species source of RG sequence, with Archaea in blue and Bacteria in red; or topoisomerase source, with Archaea in purple, Bacteria in green.
