## Supplementary material for "Positively twisted: The complex evolutionary history of Reverse Gyrase suggests a non-hyperthermophilic Last Universal Common Ancestor": SI Fig. 12

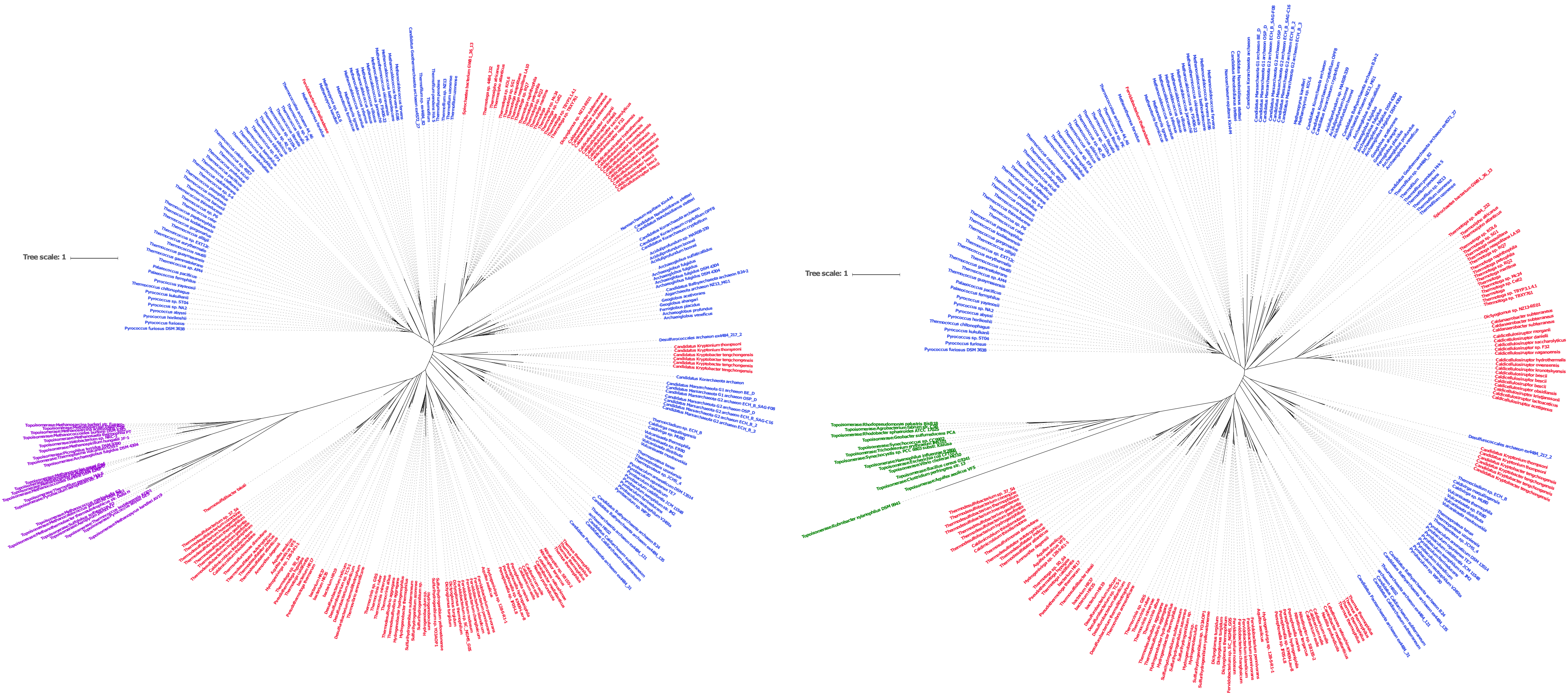
Tree scale: 1
Tree scale: 1

**SI Fig. 12.** Phylogenetic reconstruction of RG dataset with TopR1-like and TopR2-like sequences removed. Bacterial or Archaeal topoisomerase sequences included as an outgroup. Branch labels show species source of RG sequence, with Archaea in blue and Bacteria in red; or topoisomerase source, with Archaea in purple, Bacteria in green.
