## Supplementary material for "Positively twisted: The complex evolutionary history of Reverse Gyrase suggests a non-hyperthermophilic Last Universal Common Ancestor": SI Fig. 13

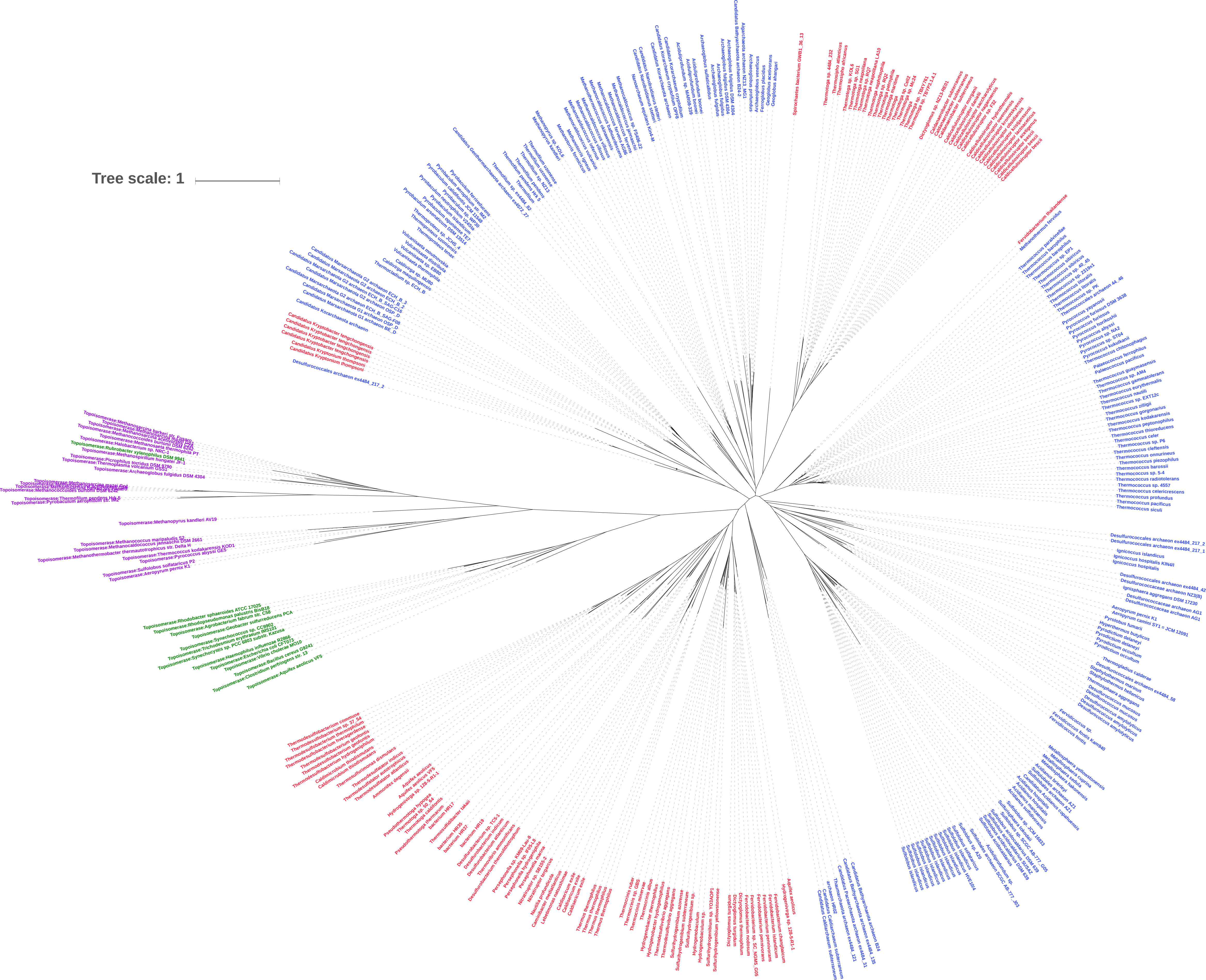
Tree scale: 1
Topoisomerase:Methanosarcina barkeri str. Fusaro
Topoisomerase:Halobacterium sp. NRC-1
Topoisomerase:Rubrobacter xylanophilus DSM 9941
Topoisomerase:Methanospirillum hungatei JF-1
Topoisomerase:Picrophilus torridus DSM 9790
Topoisomerase:Thermoplasma volcanium GSS1
Topoisomerase:Archaeoglobus fulgidus DSM 4304
Topoisomerase:Methanococcoides burtonii DSM 6242
Topoisomerase:Thermofilum pendens Hrk 5
Topoisomerase:Pyrobaculum aerophilum str. IM2
Topoisomerase:Methanopyrus kandleri AV19
Topoisomerase:Methanococcus maripaludis S2
Topoisomerase:Methanocaldococcus jannaschii DSM 2661
Topoisomerase:Methanothermobacter thermautotrophicus str. Delta H
Topoisomerase:Thermococcus kodakarensis KOD1
Topoisomerase:Pyrococcus abyssi GE5
Topoisomerase:Sulfolobus solfataricus P2
Topoisomerase:Aeropyrum pernix K1
Topoisomerase:Rhodobacter sphaeroides ATCC 17025
Topoisomerase:Rhodopseudomonas palustris BisB18
Topoisomerase:Agrobacterium fabrum str. C58
Topoisomerase:Geobacter sulfurreducens PCA
Topoisomerase:Synechococcus sp. CC9902
Topoisomerase:Trichodesmium erythraeum IMS101
Topoisomerase:Synechocystis sp. PCC 6803 substr. Kazusa
Topoisomerase:Haemophilus influenzae R2866
Topoisomerase:Escherichia coli CFT073
Topoisomerase:Vibrio cholerae MO10
Topoisomerase:Bacillus cereus G9241
Topoisomerase:Clostridium perfringens str. 13
Topoisomerase:Aquifex aeolicus VF5
Desulfurococcales archaeon ex4484_217_2
Candidatus Kryptobacter tengchongensis
Candidatus Kryptonium thompsoni
Spirochaetes bacterium GWB1_36_13
Fervidobacterium thailandense
Methanothermus fervidus
Thermococcus paralvinellae
Thermococcus barophilus
Thermococcus litoralis
Pyrococcus furiosus
Pyrococcus horikoshii
Pyrococcus abyssi
Palaeococcus ferrophilus
Palaeococcus pacificus
Thermococcus guaymasensis
Thermococcus kodakarensis
Thermococcus onnurineus
Thermococcus barossii
Thermococcus radiotolerans
Thermococcus sp. 4557
Thermococcus celericrescens
Thermococcus profundus
Thermococcus pacificus
Thermococcus siculi
Desulfurococcales archaeon ex4484_217_1
Ignicoccus islandicus
Ignicoccus hospitalis KIN4/I
Ignicoccus hospitalis
Ignisphaera aggregans DSM 17230
Aeropyrum pernix K1
Aeropyrum camini SY1 = JCM 12091
Pyrolobus fumarii
Hyperthermus butylicus
Pyrodictium delaneyi
Pyrodictium occultum
Thermogladius calderae
Staphylothermus marinus
Staphylothermus hellenicus
Thermosphaera aggregans
Desulfurococcus mucosus
Desulfurococcus amylolyticus
Fervidicoccus fontis Kam940
Fervidicoccus fontis
Metallosphaera yellowstonensis
Metallosphaera cuprina
Metallosphaera sedula
Acidianus hospitalis
Sulfolobus islandicus
Thermodesulfobacterium commune
Thermodesulfobacterium geofontis
Thermodesulfatator indicus
Thermodesulfatator atlanticus
Ammonifex degensii
Aquifex aeolicus
Hydrogenivirga sp. 128-5-R1-1
Thermus thermophilus
Thermocrinis ruber
Thermocrinis albus
Sulfurihydrogenibium azorense
Sulfurihydrogenibium yellowstonense
Dictyoglomus turgidum
Dictyoglomus thermophilum
Fervidobacterium nodosum
Fervidobacterium pennivorans
Fervidobacterium islandicum
Candidatus Bathyarchaeota archaeon B24
Candidatus Caldiarchaeum subterraneum
Thermotoga maritima
Thermotoga neapolitana
Caldicellulosiruptor bescii
Caldicellulosiruptor saccharolyticus
Thermofilum pendens
Methanopyrus kandleri
Methanocaldococcus jannaschii
Archaeoglobus fulgidus DSM 4304
Archaeoglobus veneficus
Ferroglobus placidus
Geoglobus acetivorans
Geoglobus ahangari
Nanoarchaeum equitans Kin4-M
Pyrobaculum aerophilum str. IM2
Pyrobaculum islandicum DSM 4184
Vulcanisaeta distributa
Caldivirga maquilingensis

**SI Fig. 13.** Phylogenetic reconstruction of RG dataset with TopR2-like sequences removed. Bacterial and Archaeal topoisomerase sequences included as an outgroup. Branch labels show species source of RG sequence, with Archaea in blue and Bacteria in red; or topoisomerase source, with Archaea in purple, Bacteria in green.
